# Perceptual awareness is gradual in temporal and dichotomous in fronto-parietal cortices

**DOI:** 10.1101/2022.12.14.520410

**Authors:** Marta Poyo Solanas, Minye Zhan, Beatrice de Gelder

## Abstract

Two major issues in consciousness research concern the measuring methods that determine perceptual unawareness and whether consciousness is a gradual or an ‘all-or-nothing’ phenomenon. This 7T fMRI study addresses both questions using a continuous flash suppression paradigm with an emotional recognition task (fear vs neutral bodies) in combination with the perceptual awareness scale. Behaviorally, recognition sensitivity increased linearly with increased stimuli awareness and was at chance level during perceptual unawareness. Threat expressions triggered a slower heart rate than neutral ones during ‘almost clear’ experience of the stimulus, indicating freezing behavior. The activity in occipital, temporal, parietal and frontal regions as well as in amygdala increased with increased stimulus awareness while the activity in early visual areas showed the opposite pattern. The relationship between temporal area activity and perceptual awareness was better characterized by a gradual model while the activity in fronto-parietal areas by a dichotomous model, suggesting different roles in conscious processing. Interestingly, no evidence of non-conscious processing was found in amygdala as well as no significant effect of emotion, in disagreement with the functions long ascribed to this subcortical structure.

## Introduction

Substantial evidence has been gathered in the last decades about the role of consciousness in visual perception. Yet, the assessment of subjective perceptual awareness remains a major theoretical and methodological issue in consciousness studies. For example, previous studies have often measured participants’ perceptual experience with verbal reports after the experiment, which may not be a reliable method for measuring subjective experience during a task (Pessoa, Japee, Sturman, & Ungerleider, 2006; Tsuchiya & Adolphs, 2007). Other studies have adopted more rigorous approaches through trial-by-trial assessments or by adopting an objective threshold (e.g., specific stimuli contrast after psychophysical testing), but these attempts are still susceptible to methodological biases when not formally addressed. This is the case of using percent correct values to evaluate chance performance, as one might conclude that participants are unaware of the stimuli when in fact they may still be able to reliably detect them (Green & Swets, 1966; Macmillan & Creelman, 2004). Many authors have therefore proposed the use of signal detection theory (SDT) measures (Green & Swets, 1966; Tanner & Swets, 1954) to evaluate subjective perceptual awareness independently of participants’ response bias (Green & Swets, 1966; Macmillan & Creelman, 2004). As a consequence of the efforts made to account for these issues, many studies now report no evidence of perceptual processing without accompanying awareness in healthy participants (Hedger, Adams, & Garner, 2015; Hedger, Gray, Garner, & Adams, 2016; Hoffmann, Lipka, Mothes-Lasch, Miltner, & Straube, 2012; Hoffmann, MothesLLasch, Miltner, & Straube, 2015; Mayer, Merckelbach, de Jong, & Leeuw, 1999; Pessoa, 2005; Pessoa, Japee, & Ungerleider, 2005; Straube, Dietrich, MothesLLasch, Mentzel, & Miltner, 2010).

Another source of controversy relates to the task employed to assess perceptual awareness. Early studies often used a dichotomous measure (i.e., yes/no, seen/unseen responses), which is now considered an inadequate approach for characterizing conscious perception. The reason is that such measure may not capture intermediate states of experience and thus may not correctly differentiate genuine forms of blindsight from residual conscious vision (Mazzi, Bagattini, & Savazzi, 2016). This view has led to the development of finer measures of perceptual awareness, such as the perceptual awareness scale (PAS), with four different response alternatives: ‘no experience’, ‘brief glimpse’, ‘almost clear experience’ and ‘clear experience’ (Ramsøy & Overgaard, 2004). Recent studies using PAS have reported chance performance during perceptual unawareness in objective forced-choice discrimination tasks, as well as intermediate states of perceptual awareness between unseen and completely seen reports (Hesselmann, Darcy, Rothkirch, & Sterzer, 2018; Lähteenmäki, Hyönä, Koivisto, & Nummenmaa, 2015; Lamy, Alon, Carmel, & Shalev, 2015; Lamy, Carmel, & Peremen, 2017; Lohse & Overgaard, 2019; Peremen & Lamy, 2014; Ramsøy & Overgaard, 2004; Tagliabue, Mazzi, Bagattini, & Savazzi, 2016). These findings have therefore instigated debates about non-conscious processing but also sparked theoretical discussions about whether perceptual awareness is a graded or an ‘all-or-none’ phenomenon. Despite all the research efforts trying to solve this controversy, a consensus has not yet been achieved as both views are supported by strong empirical evidence (for a review see Windey & Cleeremans, 2015).

In the long standing debate about perception without awareness as well as in dichotomous vs. gradual discussions on consciousness, emotional stimuli have not occupied a major place. Indeed, in the dominant theories about consciousness, such as global workspace theories, higher order theories, integrated information theory or re-entry and predictive processing theories, the debates mainly concern cognitive processes (Seth & Bayne, 2022). It is therefore an open question whether those major theories and debates on consciousness are applicable to affective stimuli. In this regard, fearful stimuli are considered a particularly strong candidate for non-conscious processing (de Gelder, Morris, & Dolan, 2005; Morris, de Gelder, Weiskrantz, & Dolan, 2001; Vieira, Wen, Oliver, & Mitchell, 2017; Whalen et al., 1998) and appear to gain privileged access to awareness in comparison to other emotions (Gray, Adams, Hedger, Newton, & Garner, 2013; Lee, Lim, Lee, & Choi, 2009; Oliver, Mao, & Mitchell, 2015; Stein, Seymour, Hebart, & Sterzer, 2014; Tsuchiya, Moradi, Felsen, Yamazaki, & Adolphs, 2009; Yang, Zald, & Blake, 2007). Yet, most research on perceptual awareness and affective perception has used facial expressions. Available research on affective non-conscious processing has revealed differences between facial and bodily expressions even if they presumably represent the same emotion. For example, Zhan and colleagues (2015) found that angry bodies had shorter suppression times in comparison to other bodily emotions, while angry facial expressions had the longest suppression times (Zhan, Hortensius, & de Gelder, 2015). These findings evidence the importance of extending investigations to other sources of affective information.

With regards to brain processes, a subcortical pathway involving the superior colliculus, pulvinar and amygdala has been suggested to be crucial in the non-conscious processing of emotional expressions (Tamietto & de Gelder, 2010). Amygdala is known to have a key role in affective processing (Davis & Whalen, 2001), while pulvinar has been involved in visual attention, saliency detection, spatial processing and top-down anticipation of visual information (Saalmann & Kastner, 2011). In healthy participants, previous masking (Morris, Öhman, & Dolan, 1998; Whalen et al., 1998) and binocular rivalry studies (Pasley, Mayes, & Schultz, 2004; Williams, Morris, McGlone, Abbott, & Mattingley, 2004) have reported amygdala responses to stimuli in conditions of unawareness. Yet again, evidence of amygdala involvement in non-conscious processing of affective information mainly comes from face studies using dichotomous measures. Emotional bodies have also shown to activate amygdala (Hadjikhani & de Gelder, 2003; Kret, Pichon, Grèzes, & de Gelder, 2011; Sinke, Sorger, Goebel, & de Gelder, 2010; van de Riet, Grèzes, & de Gelder, 2009), but their processing outside conscious awareness has yielded mixed results for this subcortical structure (de Gelder & Hadjikhani, 2006; Tamietto et al., 2015; Van den Stock et al., 2011; Van den Stock et al., 2014; Zhan, Goebel, & de Gelder, 2018). With regards to cortical areas, previous work has suggested a crucial role of the frontal, parietal and temporal cortex in perceptual stimulus awareness (Kreiman, Fried, & Koch, 2002; Leopold & Logothetis, 1996; Logothetis & Schall, 1989; Panagiotaropoulos, Deco, Kapoor, & Logothetis, 2012; Sheinberg & Logothetis, 1997). However, evidence for the involvement of these areas in the non-conscious processing of affective stimuli is still limited.

It is therefore an open question whether affective signals, especially body expressions, are processed under conditions of perceptual unawareness and whether the perception of body expressions presents a gradual or a dichotomous relationship to perceptual awareness. Here, we used the CFS paradigm and ultrahigh field 7T (f)MRI scanning to investigate the processing of threat stimuli (fearful vs. neutral body expressions) in healthy participants. Body expressions were randomly presented either to the left or right visual field of participants’ non-dominant eye, while a colorful dynamic noise mask was shown to the dominant eye (**Figure 1**). Participants performed a two-alternative forced-choice task (fear/neutral) followed by the rating of their visual stimulus experience with the perceptual awareness scale (Ramsøy & Overgaard, 2004). Our experiment has several improvements over previous studies. First, we used the CFS paradigm (Tsuchiya & Koch, 2005) to render stimuli invisible. This method has been increasingly used since it creates a stronger suppression and more stable non-conscious perception in comparison to previous methods, such as masking (Yang, Brascamp, Kang, & Blake, 2014). Second, we measured perceptual awareness on a trial-by-trial basis and by using the perceptual awareness scale. This allowed us to differentiate genuine forms of perception without awareness from conditions of partial perceptual awareness and also allowed us to assess whether perceptual awareness is a gradual or a dichotomous phenomenon. Importantly, subjective perceptual awareness reports were also formally and objectively assessed with signal detection theory measures (Green & Swets, 1966) and by comparing task performance to chance level. In addition, with ultra-high field magnetic resonance imaging we were able to obtain higher spatial resolution and specificity but also a higher contrast-to-noise ratio (Uğurbil, 2014). Finally, the use of body expressions provided a novel take on the processing of social information beyond facial expressions.

**Figure 1.**
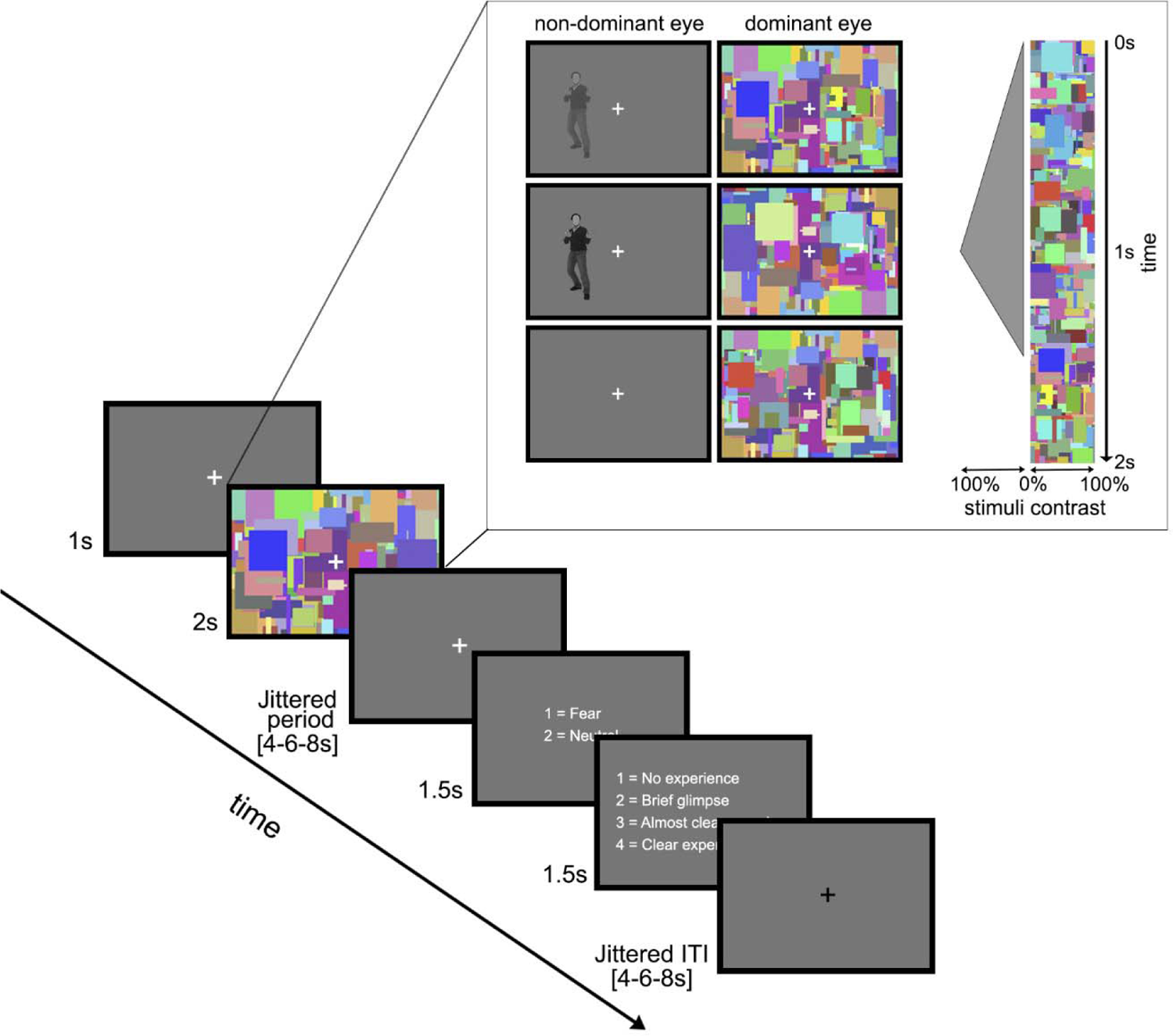
Schematic view of a trial presentation sequence in the main experiment. After a 1s-fixation period, a 2s-CFS period started with the gradual ramping up of the body stimulus contrast from 0% to full contrast over 1s, followed by the contrast reduction to 0% within 0.5s and a 0.5s blank period (see content within frame). The contrast of the dynamic colorful mask was constant throughout the 2s. However, both the body stimuli and the mask contrasts were determined for each trial using a staircase procedure with 10 steps (body stimuli: 5%, 14%, 23%, 32%, 41%, 50%, 50%, 50%, 50%, 50%; noise: 100%, 100%, 100%, 100%, 100%, 82%, 64%, 46%, 28%, 10%) that depended on the participant’s visual experience of the stimulus in the previous trial. After a jittered fixation period (4-6-8s), participants were required to make two active responses, each within a 1.5s window: a two-alternative forced-choice task (fear vs. neutral) and the rating of their visual experience of the stimulus according to the Perceptual Awareness Scale (PAS). The inter-trial-interval (ITI) was jittered (4-6-8s) and the average trial duration was 18s.

## Results

### Behavioral results

#### Sensitivity

The analysis of recognition sensitivity showed a significant main effect of Perceptual Awareness (F (3, 41.76) = 37.13, p < .001), indicating a significantly higher sensitivity during ‘brief glimpse’ (PAS2, M = 0.38, SE = 0.08), ‘almost clear’ (PAS3, M = 1.32, SE = 0.13) and ‘clear’ (PAS4, M = 1.36, SE = 0.18) experience conditions than during ‘no experience’ (PAS1, M = −0.03, SE = 0.03). Sensitivity was also significantly lower during ‘brief glimpse’ than for ‘clear’ and ‘almost clear’ experience (see **Figure 2A**). Sensitivity values differed from the chance level in PAS2 to PAS4 (p < .001), but not in PAS1 (p = .352). Further analyses indicated that the relationship between perceptual sensitivity and perceptual awareness was significantly better described by a gradual model (M = 1.05, SE = 1.61) than by a dichotomous one (M = 6.61, SE = 1.13; t(16) = −3.69, p = .002).

**Figure 2.**
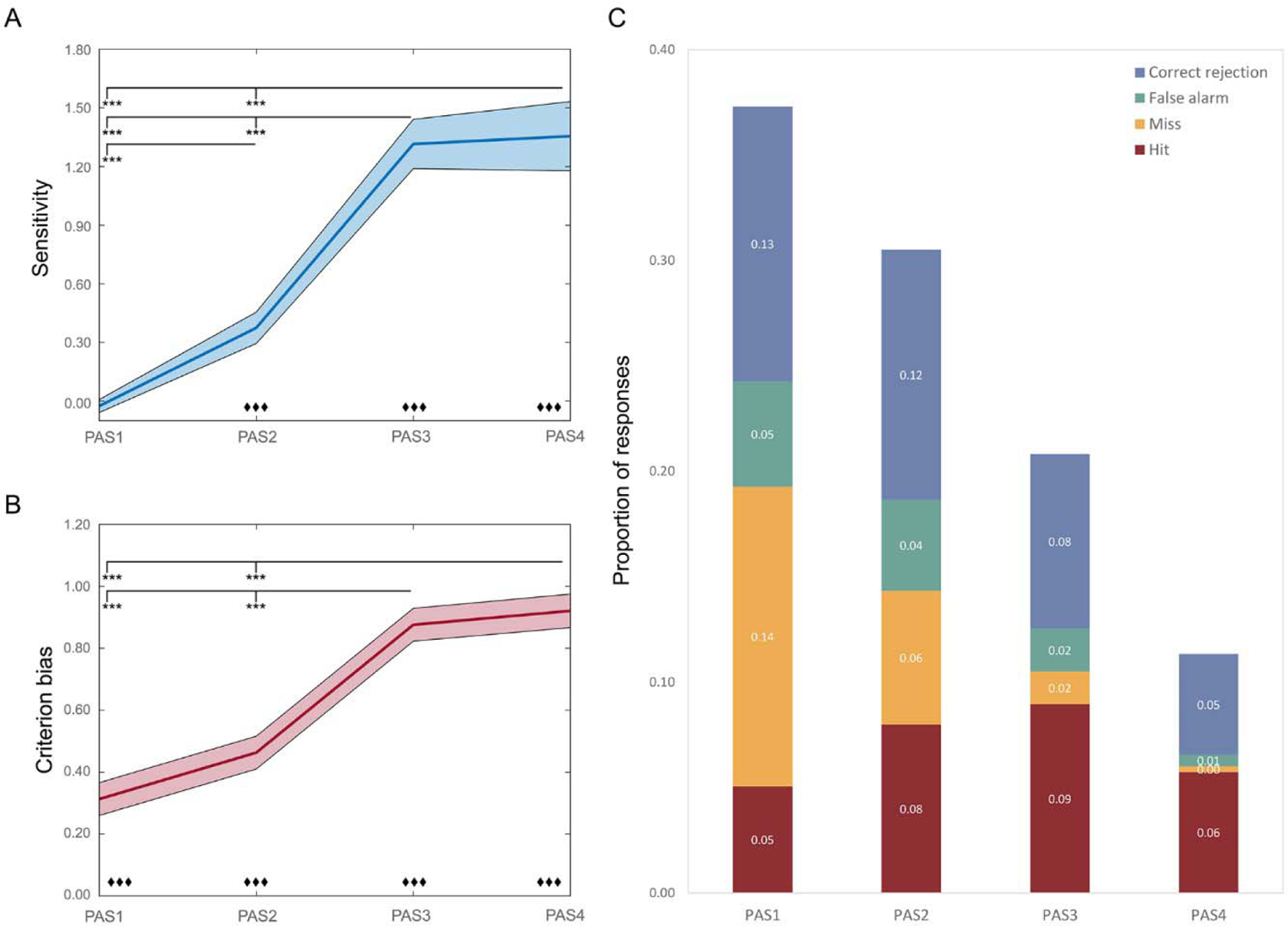
Overview of behavioral results. **A)** Estimated marginal means of sensitivity values separated by PAS levels. Shadowed area indicates standard error from the mean; **B)** Estimated marginal means of criterion bias values separated by PAS levels. Shadowed area indicates standard error from the mean; **C)** Average proportion of responses separated by PAS and SDT measures (i.e., Hit, Miss, False alarm, Correct rejection) (see *Supplementary Table S2* for average values and standard errors). *Hit*: correctly categorized fearful stimuli; *Miss*: fearful stimuli categorized as neutral; *False alarms*: neutral stimuli categorized as fearful; *Correct rejection*: correctly categorized neutral stimuli. ***Notes***: Black asterisks denote significant differences between PAS levels. Rhombi denote significant difference from zero. */♦: p < .05; **/♦♦: p < .01; ***/♦♦♦: p < .001. ***Abbreviations***: PAS: perceptual awareness scale: PAS1: ‘no experience’, PAS2: ‘brief glimpse’, PAS3: ‘almost clear experience’, PAS4: ‘clear experience’.

#### Criterion bias

The criterion bias analysis showed a significant main effect of Perceptual Awareness (F(3, 23.17) = 26.01, p < .001) with more conservative emotional categorizations for ‘clear’ (M = 0.93, SE = 0.05) and ‘almost clear’ (M = 0.88, SE = 0.05) experiences than ‘no experience’ (M = 0.31, SE = 0.05) or ‘brief glimpse’ (M = 0.46, SE = 0.05) (see **Figure 2B**). Criterion bias values did not differ significantly between ‘clear’ and ‘almost clear’ experiences as well as between ‘no experience’ and ‘brief glimpse’. Criterion bias scores were different from zero at all levels of the perceptual awareness scale (p < .001).

#### Reaction times

The analysis of RTs of the emotional categorization task (four levels: hit, miss, false alarm and correct rejection) did not show differences between different response types (F(3, 27.62) = 0.78, p = .513) (see **Table S1** in *Supplementary Information*). The analysis of the RTs of the perceptual awareness ratings (four levels: PAS 1-4) showed a main effect of Perceptual Awareness (F (3, 21.77) = 5.29, p = .007), indicating significant faster responses for ‘no experience’ (M = 0.48, SE = 0.04) than for ‘brief glimpse’ (M = 0.56, SE = 0.04) and marginally faster than for ‘clear’ ratings (M = 0.58, SE = 0.04) (see **Table S1** in *Supplementary Information*). For an overview of the proportion of responses by Signal Detection Theory measure and Perceptual Awareness Scale rating, see **Table S2** in *Supplementary Information*.

### Brain results

#### Regions sensitive to perceptual awareness

A group ANOVA with within-subjects factor Perceptual Awareness (i.e., PAS1-4) was performed to localize the areas sensitive to different degrees of perceptual awareness. ROIs showing a main effect of perceptual awareness were found bilaterally in the lateral occipito-temporal cortex (LOTC), fusiform gyrus, inferior temporal gyrus (ITG) as well as in right amygdala and left precuneus, occipital cortex, intraparietal sulcus (IPS), inferior frontal cortex (IFC), superior temporal gyrus (STG) and posterior superior temporal sulcus (pSTS) (see **Figure 3**; see Supplementary **Table S3** for more details on ROI location, size and statistical values).

**Figure 3.**
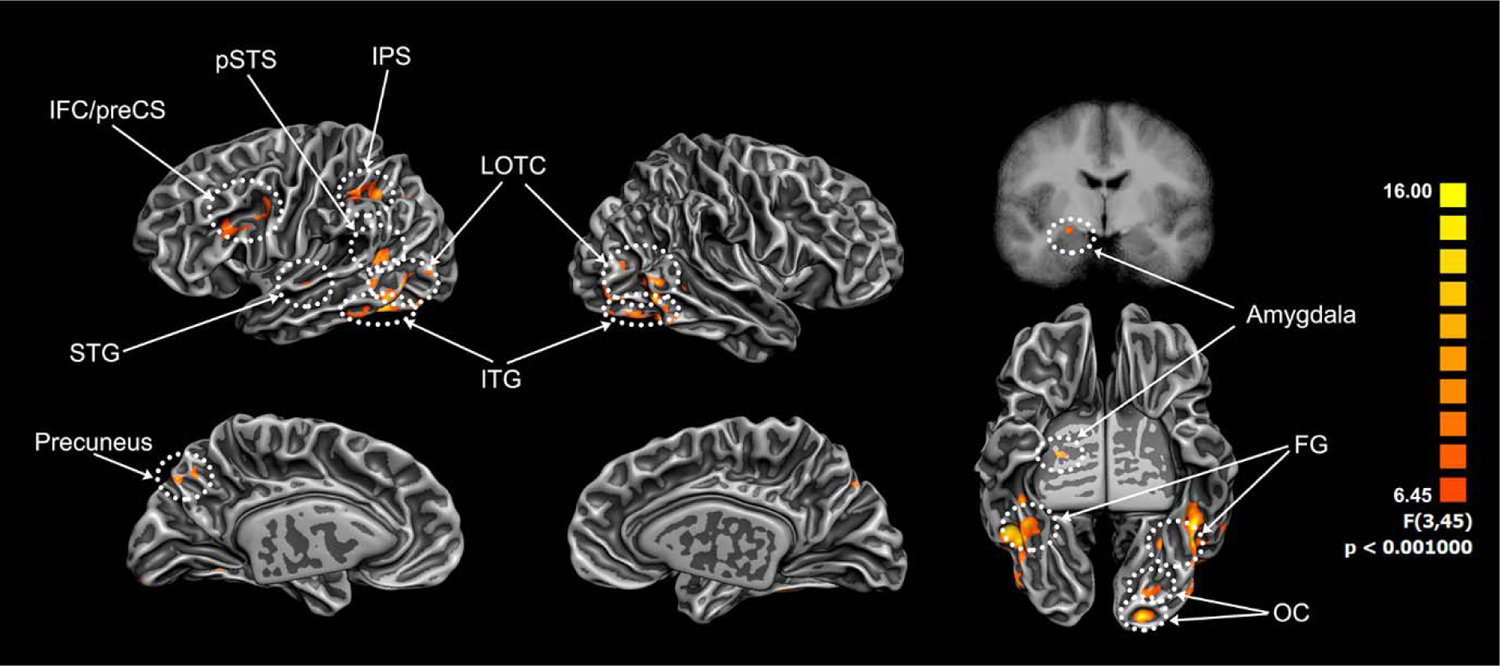
Areas showing a main effect of perceptual awareness at the group level. These ROIs resulted from the repeated measures ANOVA (N = 16) with within-subject factor Perceptual Awareness (four levels: PAS1-4) (cluster size corrected with Monte-Carlo simulation, alpha level = 0.05, initial p = 0.001, numbers of iterations = 5000). *Abbreviations*: **FG**: fusiform gyrus; **IFC**: inferior frontal cortex **IPS**: intraparietal sulcus; **ITG**: inferior temporal gyrus; **LOTC**: lateral occipito-temporal cortex; **OG**: occipital gyrus; **preCS**: precentral sulcus; **pSTS**: posterior superior temporal sulcus; **STG**: superior temporal gyrus.

#### Effect of emotion and perceptual awareness on PAS-sensitive ROI activity

To further understand their involvement in perceptual awareness, the beta values of these ROIs were then analyzed with a linear mixed-model analysis with within-subject factors Emotion (two levels: Neutral & fear) and Perceptual Awareness (four levels: PAS1-4). All the ROIs showed a significant main effect of PAS but no significant main effect of Emotion or interaction effect (**Table 1**; see Supplementary **Table S4** for further statistical details; see **Figure 4** for ROI activity visualization). Subsequently, different pairwise comparisons were performed to understand differences in activity levels between non-conscious and conscious perception (PAS1 vs PAS4; PAS1 vs PAS3), between non-conscious and threshold vision (PAS1 vs PAS2), between threshold and clearer degrees of conscious perception (PAS2 vs PAS4; PAS2 vs PAS3) and to assess whether there was a difference between levels of conscious perceptual awareness (PAS3 vs PSA4) in the different ROIs.

**Figure 4.**
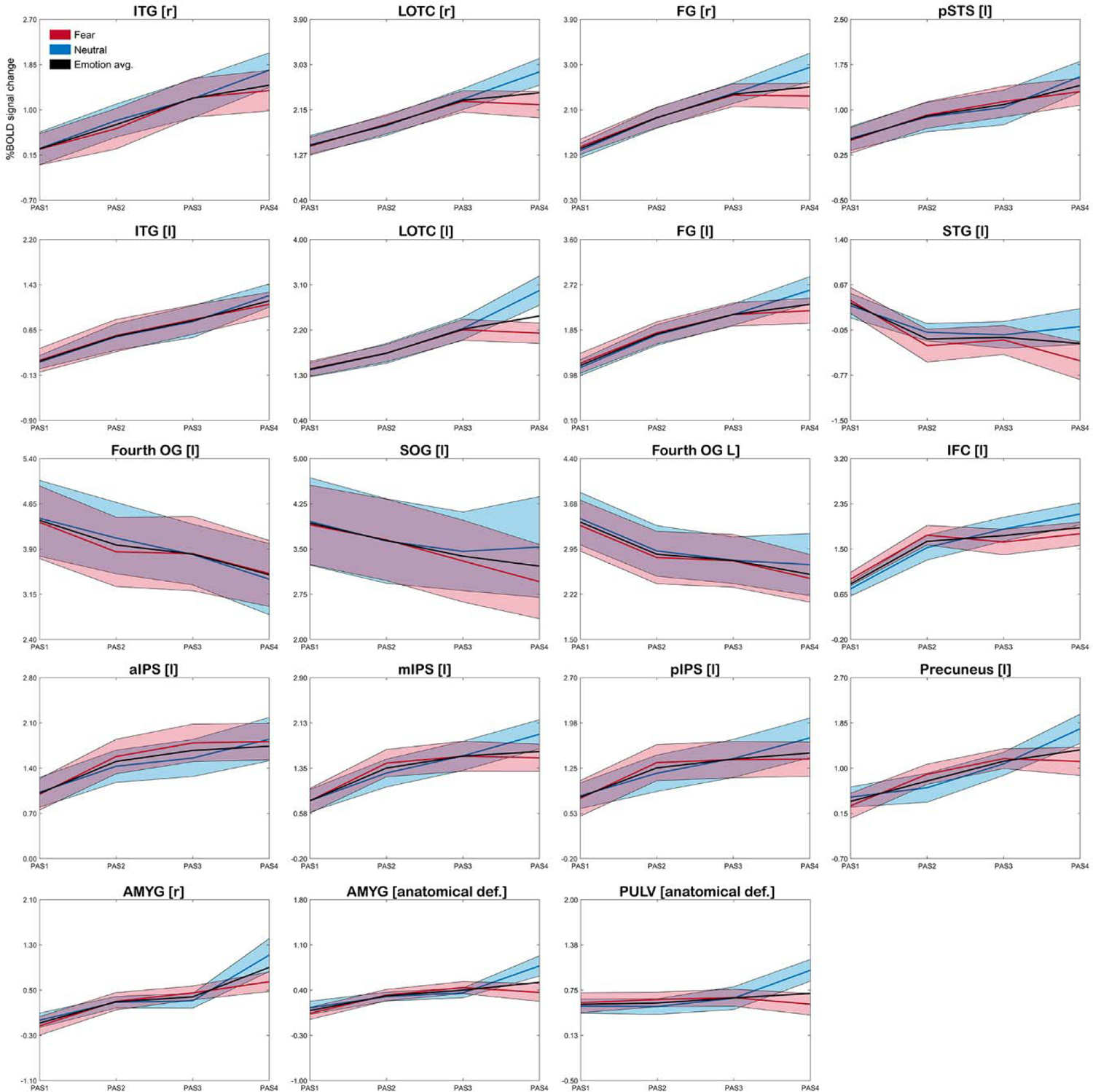
Brain responses across Perceptual Awareness levels separately for each stimuli type. Brain responses represent %-BOLD signal changes of the ROIs showing a main effect of PAS as well as the anatomically defined amygdala and pulvinar. Shadowed area indicates standard error from the mean. *Abbreviations*: **aIPS**: anterior intraparietal sulcus; **def**.: definition; **FG**: fusiform gyrus; **Fourth OG**: fourth occipital gyrus; **IFC**: inferior frontal cortex; **ITG**: inferior temporal gyrus; **LOTC**: lateral occipito-temporal cortex; **mIPS**: medial intraparietal sulcus; **PAS**: perceptual awareness scale; **pIPS**: posterior intraparietal sulcus; **preCS**: precentral sulcus; **pSTS**: posterior superior temporal sulcus; **SOG**: superior occipital gyrus; **STG**: superior temporal gyrus. **[l]**: left hemisphere; **[r]**: right hemisphere.

**Table 1.**
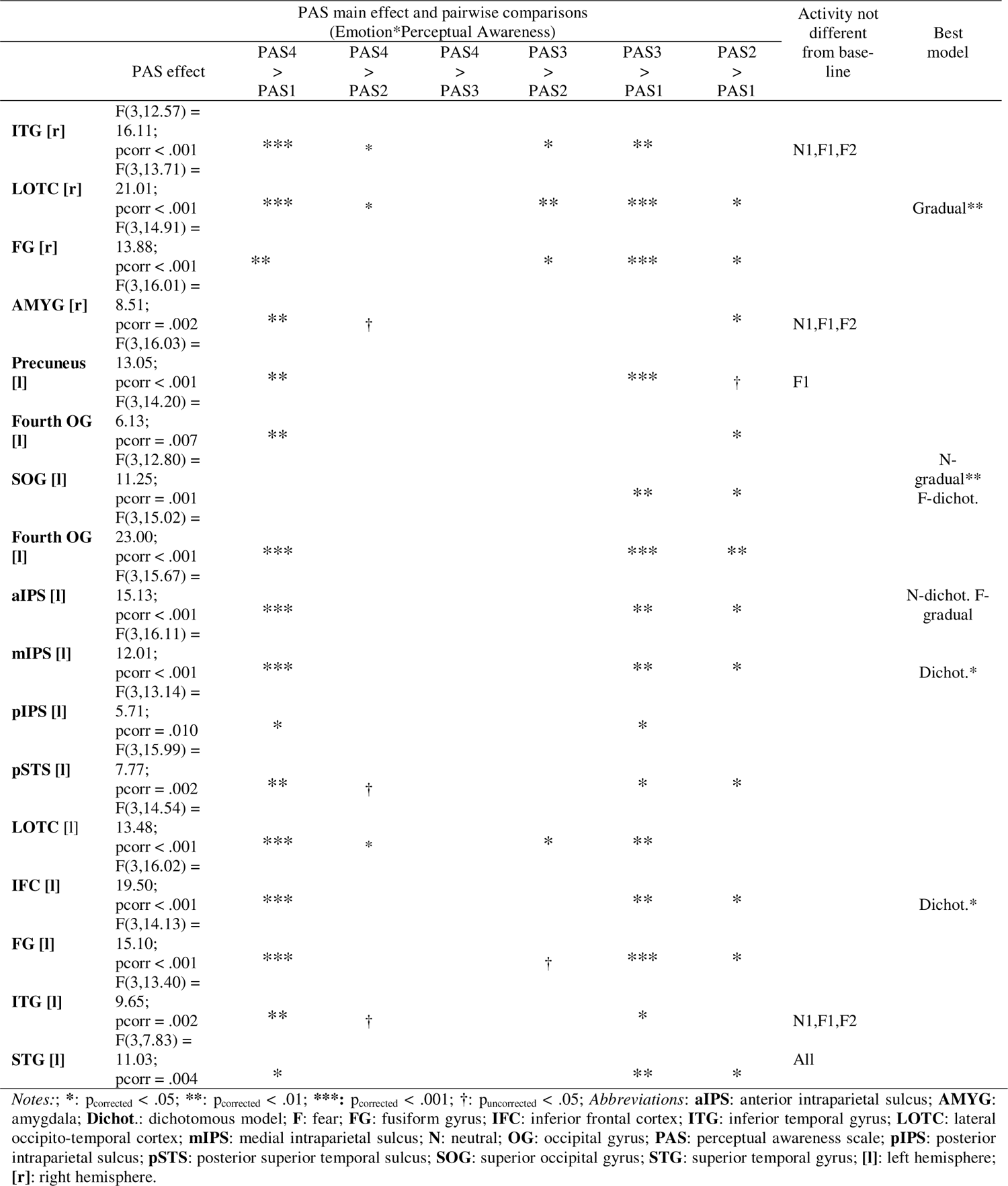
Results of the linear mixed model analysis with within-subject factors Emotion and PAS and the model fitting, separately for each ROI.

Significantly higher activity for PAS4 than PAS1 was observed in all ROIs with the exception of the left superior occipital gyrus (SOG), left STG and two areas in the left fourth occipital gyrus (see **Table 1**). Overall, similar results were observed for the PAS3 and PAS1 comparison. The ROIs in the left fourth occipital gyrus, left STG and SOG showed the opposite pattern, with higher activity for PAS1 than PAS3 and PAS4 (**Figure 4**), although only significantly for the former two ROIs (**Table 1**). Significantly higher activity for PAS4 in comparison to PAS2 was found in bilateral LOTC and ITG, right amygdala and left pSTS, although not surviving multiple comparisons correction. Pairwise comparisons revealed that the activity for PAS3 and PAS4 conditions was not significantly different in any of the defined ROIs. Most of the ROIs showed a significantly different activity level between PAS2 and PAS1 conditions, except the bilateral ITG, left IPS and the left LOTC.

Overall, most of the defined ROIs showed an activity level significantly different from zero across the eight Emotion*Perceptual Awareness conditions (**Table 1**; see Supplementary **Table S5** for further statistical details; see **Figure 4** for ROI activity visualization). An exception was found in the right amygdala and bilateral ITG, where the activity level was not significantly different from zero at PAS1 for both emotions, and at PAS2 for fear. In addition, the activity in the left precuneus did not differ from zero at PAS1 for fearful body expressions. The activity in the left STG did not differ from zero at any of the Emotion*Perceptual Awareness conditions.

To further investigate whether perceptual awareness is a gradual or a dichotomous phenomenon, two linear mixed models corresponding to each phenomenon were fit to the data. The activity pattern in the right LOTC was significantly better described by a gradual model, regardless of the emotion of the stimuli (**Table 1**; see Supplementary **Table S6** for further statistical details; see Supplementary **Table S7** for results on model slope and intercept comparisons). The responses in the left medial IPS and IFC were significantly best described by a dichotomous model, also with no differences across emotions. Yet, emotion specificity was found in some areas. The activity pattern in the SOG was different for the neutral and fearful body expressions across all PAS levels (i.e., significant interaction effect). For neutral bodies, SOG activity was significantly best described by a gradual model while in the case of fearful bodies, the activity was better represented by a dichotomous model, although it did not reach significance in the latter case. The responses of the left anterior IPS also showed an Emotion*Perceptual Awareness interaction effect, with the activity elicited by neutral bodies better described by a dichotomous model while a gradual model better described the activity elicited by fearful body expressions. However, pairwise comparisons were not significant in both cases. The rest of the ROIs did not show a significant preference for either of the models.

#### Effect of emotion and perceptual awareness in anatomically defined amygdala and pulvinar

The analysis of amygdala betas yielded a significant main effect of Perceptual Awareness (F(3, 14.39) = 8.60, p = .002), showing that amygdala activity was significantly lower during ‘no experience’ (M = 0.08, SE = 0.09) than during ‘brief glimpse’ (M = 0.31, SE = 0.06), ‘almost clear’ (M = 0.40, SE = 0.07) and ‘clear experience’ (M = 0.58, SE = 0.13) (see **Figure 4**). This analysis also yielded a significant main effect of Emotion (F(1, 13.23) = 6.67, p = .023), indicating a higher amygdala activity for neutral (M = 0.38, SE = 0.07) than fearful body expressions (M = 0.30, SE = 0.07). One-sample t-tests revealed that amygdala activity differed from baseline at PAS2 to PAS4 (p < .001) but not at PAS1 (i.e., ‘no experience’), for both emotions. Amygdala activity did not show a significant preference for either the gradual or the dichotomous model.

The analysis of the activity in pulvinar did not show significant main effects for Emotion or Perceptual Awareness nor a significant Emotion*Perceptual Awareness interaction (see **Figure 4**). However, pulvinar responses were significantly different from zero in all PAS levels for both emotions (p < .005). As for the amygdala, pulvinar did not show a significant preference for either the gradual or the dichotomous model.

### Heart rate

The analysis of heart rate responses yielded a marginal significant effect of Perceptual Awareness (F(3, 106.30) = 2.68, p = .051) and a significant Emotion*Perceptual Awareness interaction (F(3, 106.03) = 2.75, p = .047), indicating that at PAS3, heart rate in response to fearful bodies (M = −3.59, SE = 0.62) was slower than to neutral bodies (M = −1.91, SE = 0.63). For fearful expressions, heart rate was marginally significantly higher during ‘no experience’ (M = −1.54, SE = 0.63) and ‘clear’ experience (M = −1.50, SE = 0.63) than during ‘almost clear’ experience (M = −3.59, SE = 0.62) (see **Figure 5**).

**Figure 5.**
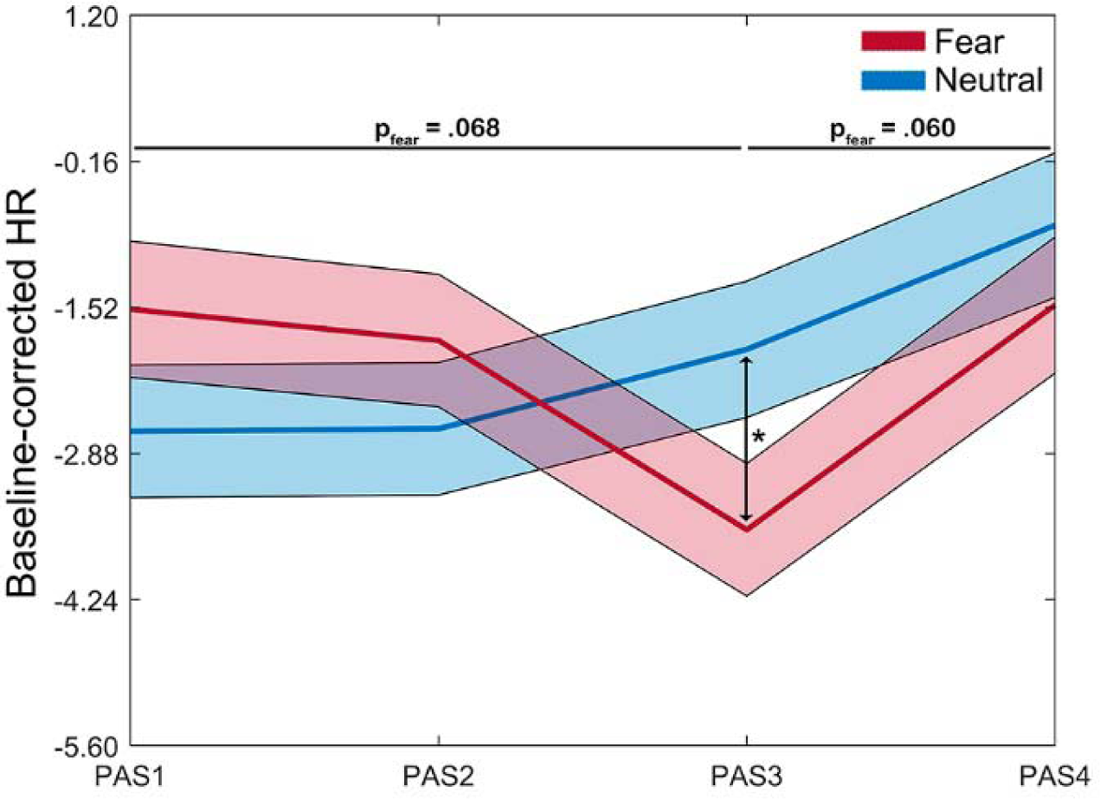
Heart rate across perceptual awareness levels and emotional stimuli categories. Heart rate (beats per minute) was baseline-corrected and averaged over the 2s-CFS period. Shadowed area indicates standard error from the mean. *Abbreviations*: **\***: p < .05; **HR**: heart rate; **PAS**: perceptual awareness scale.

## Discussion

This ultra-high field fMRI study investigated the perception of threat stimuli (fearful vs. neutral body expressions) in healthy participants using the continuous flash suppression paradigm in combination with the perceptual awareness scale. Behaviorally, we found a gradual relationship between recognition sensitivity and perceptual awareness and no evidence of perceptual discrimination without perceptual awareness. Heart rate was slower for fearful than neutral bodies during almost clear stimulus perception, in line with freezing behavior. At the brain level, the activity in occipital, temporal, parietal and frontal regions as well as in amygdala increased with increased stimulus awareness, while the activity in early visual areas showed the opposite pattern. The relationship between temporal cortex activity and perceptual awareness was better characterized by a gradual model while the activity in fronto-parietal areas by a dichotomous model, suggesting different roles in conscious processing. Interestingly, no evidence of non-conscious body expression processing was found in the amygdala as well as no increased activation for threat stimuli, in contrast to some findings in the literature (Adolphs, Tranel, Damasio, & Damasio, 1995; Tamietto & de Gelder, 2010).

### Behavioral evidence for the gradual account but not for non-conscious processing

The behavioral results revealed a continuum of intermediate states of perceptual awareness (**Figure 2C**) as well as a linear increase in recognition sensitivity with increased perceptual awareness (**Figure 2A**). In line with our findings, support for the gradual view has been reported for the perception of low-level features such as color or shape (Lähteenmäki et al., 2015; Overgaard, Rote, Mouridsen, & Ramsøy, 2006; Overgaard & Sandberg, 2012; Ramsøy & Overgaard, 2004; Sandberg & Overgaard, 2015; Sandberg, Timmermans, Overgaard, & Cleeremans, 2010; Wierzchoń, Paulewicz, Asanowicz, Timmermans, & Cleeremans, 2014; Windey, Vermeiren, Atas, & Cleeremans, 2014), but also for high-level object and semantic (e.g., emotion) perception (Lähteenmäki et al., 2015; Lohse & Overgaard, 2019; Poyo Solanas, Zhan, & de Gelder, 2022). Moreover, our behavioral results replicated our earlier study using identical fearful and neutral body stimuli as well as angry body expressions (Poyo Solanas et al., 2022).

In contrast with previous work reporting non-conscious processing of emotional information in healthy participants (e.g., Khalid & Ansorge, 2017; Vieira et al., 2017; Watanabe & Haruno, 2015), we found no behavioral evidence for body expression discrimination during perceptual unawareness (**Figure 2A**). One possible reason for these divergent results may be that most studies reporting non-conscious affective processing used facial expressions (Tamietto & de Gelder, 2010). In this regard, there is evidence suggesting that emotional faces and bodies may be processed differently during CFS (Zhan et al., 2015). Another possible reason for this discrepancy may be due to methodological differences regarding the assessment of perceptual awareness. Earlier studies mostly relied on dichotomous measures, which may not be adequate for correctly differentiating true perceptual unawareness from partial/degraded states of perceptual awareness (e.g., ‘brief glimpse’ in PAS) (Mazzi et al., 2016). In agreement with our findings, growing evidence fails to find evidence for emotion processing outside conscious awareness when using finer scales of perceptual awareness in combination with objective force-choice discrimination tasks to assess task performance (Hesselmann et al., 2018; Lähteenmäki et al., 2015; Lamy et al., 2015; Lamy et al., 2017; Lohse & Overgaard, 2019; Peremen & Lamy, 2014; Poyo Solanas et al., 2022; Ramsøy & Overgaard, 2004; Tagliabue et al., 2016) as well as when controlling for other methodological confounds (Hedger et al., 2015; Peters et al., 2017; Peters & Lau, 2015; Rajananda, Zhu, & Peters, 2020).

### Different brain areas involved in gradual and ‘all-or none’ perceptual awareness

Several areas spanning early visual as well as temporal, parietal and frontal regions were influenced by perceptual awareness. Among the areas found bilaterally, there was the LOTC (overlapping with the extrastriate body area (EBA), the fusiform gyrus (overlapping with the fusiform body area (FBA), and the inferior temporal cortex, all high-level visual object areas known to be involved in object recognition (Downing, Jiang, Shuman, & Kanwisher, 2001; Peelen & Downing, 2005) (**Figure 3**). The activity in these areas increased with increased stimulus sensitivity, supporting the notion that degrees of perceptual (object) awareness correspond to degrees of stimulus recognition. Fusiform activity modulations have previously been reported in response to participants’ confidence in recognizing an object (Bar et al., 2001) as well as in response to stimulus visibility (Pessoa et al., 2006). In a study using an object naming task, recognition performance correlated with the activity of object areas in the occipito-temporal and fusiform cortex when stimuli exposure duration was varied from 20 to 500 milliseconds (Grill-Spector, Kushnir, Hendler, & Malach, 2000). Another study using body postures and a CFS paradigm found higher activation in ventral body-sensitive areas during visible conditions, although this study did not assess perceptual awareness on a trial-by-trail basis (Zhan et al., 2018). Taken together, these studies and the current findings clearly demonstrate a correlation between the level of perceptual awareness and the magnitude of activation of high-level visual areas, including body-selective ones. This picture is different from the proposal by Windey and Cleeremans (2015) that high-level visual perception is dichotomous while low-level visual processing is graded (Windey & Cleeremans, 2015). However, the recent study by Poyo Solanas and colleagues (2020) showed that a region in the lateral occipito-temporal cortex overlapping with the one of the current study encodes the degree of limb contraction. The gradual pattern found in this area may therefore reflect the encoding of this postural midlevel body feature (de Gelder & Poyo Solanas, 2021; Poyo Solanas et al., 2020) instead of a high-level body representation.

Furthermore, the current results show an effect of perceptual awareness along the intraparietal sulcus. This region is part of the dorsal attention network and responds to stimulus salience, direction of attention and saccades (Corbetta & Shulman, 2011), all of which may have played a role in our experiment. It is worth noting that our stimuli were presented at either the left or right of the fixation cross (**Figure 1**). Although participants were specifically instructed to always fixate at the cross and to refrain to do any eye movements when perceiving something in the noise, we cannot exclude that stimulus visibility may have led to saccades (Rothkirch, Stein, Sekutowicz, & Sterzer, 2012). However, in our previous pupillometry study using a similar experimental design, participants maintained central fixation in 95% of the trials (Poyo Solanas et al., 2022), which makes eye movements an unlikely explanation for the perceptual awareness modulations found in IPS. In addition, previous studies have shown increased activity in IPS for visible tools (Hesselmann, Hebart, & Malach, 2011) and inverted bodies (Zhan et al., 2018) in comparison to their unseen counterparts. Transcranial magnetic stimulation (TMS) over IPS has also shown to cause perceptual vanishing of visual stimuli (Brascamp, Sterzer, Blake, & Knapen, 2018; Kanai, Muggleton, & Walsh, 2008), suggesting that this area may mediate perception of stimuli entering consciousness.

In line with previous work, our results also suggest an important role of the inferior frontal cortex in perceptual awareness (**Figure 3**). For example, neuroimaging studies investigating bistable perception have reported increases in prefrontal activity to perceptual changes regardless of stimulus-driven transitions or of the stimuli or paradigm employed (Brascamp et al., 2018; Lumer, Friston, & Rees, 1998; Sterzer & Kleinschmidt, 2007; Zaretskaya, Thielscher, Logothetis, & Bartels, 2010). Moreover, atypical perceptual transitions during bistable perception have been observed in patients with prefrontal cortex lesions (Meenan & Miller, 1994; Ricci & Blundo, 1990; Windmann, Wehrmann, Calabrese, & Güntürkün, 2006). However, it has been claimed that the activity in the prefrontal cortex may reflect participants’ reports instead of actual changes in conscious perception (Koch, Massimini, Boly, & Tononi, 2016; Tsuchiya, Wilke, Frässle, & Lamme, 2015), which would impact the interpretability of the current results. Indeed, neuroimaging (Lumer & Rees, 1999) as well as intracranial electrophysiological (Noy et al., 2015) studies involving no-report paradigms found no involvement of the dorsolateral prefrontal cortex in perceptual awareness. Yet, activation was still observed in the IFC, the area of the pre-frontal cortex observed in the current study. Importantly, a causal role of the IFC in conscious experience has been recently demonstrated in a study combining neuroimaging methods with TMS showing that this area facilitates changes in conscious experience by continually monitoring conscious representations and comparing them to available sensory data (Weilnhammer et al., 2021).

Taken together, the current findings build upon previous work suggesting that the fronto-parietal and temporal cortex constitute a cortico-cortical network involved in perceptual stimulus awareness (Kreiman et al., 2002; Leopold & Logothetis, 1996; Logothetis & Schall, 1989; Panagiotaropoulos et al., 2012; Sheinberg & Logothetis, 1997). On one side, the gradual relationship observed between perceptual awareness and temporal cortex activity (**Table 1**) suggests that this area may encode subjective stimuli perception rather than merely physical stimuli properties (body stimuli were always presented and in the same manner). On the other side, the dichotomous activity pattern observed in anterior and medial parts of the IPS as well as in IFC suggests that these areas may be crucial in mediating stimuli entering into consciousness (Brascamp et al., 2018; Kanai et al., 2008; Weilnhammer et al., 2021). This was observed as a significantly lower activity level during ‘no experience’ (when no perceptual switches occurred) than in the rest of perceptual awareness levels (when perceptual switches occurred) as well as a non-significant activity difference between the PAS levels in which perceptual switches occurred (i.e., PAS2-4) (see **Table 1**). The increase in fronto-parietal activity with increased perceptual awareness was seen together with a decrease in the activity of early visual areas (**Figure 4**). This could reflect the increased inhibition of neural populations representing the dominant stimulus (flickering colorful mask) as the body stimulus gradually became subjectively more visible (Gail, Brinksmeyer, & Eckhorn, 2004; Keliris, Logothetis, & Tolias, 2010; Leopold & Logothetis, 1996; Maier, Logothetis, & Leopold, 2007; Wilke, Logothetis, & Leopold, 2006), suggesting that early visual areas may be sufficient to determine the content of perception in situations in which perceptual conflict has not escalated (i.e., PAS1) (Xu et al., 2016). This is in line with previous work suggesting that fronto-parietal areas may detect perceptual conflict via bottom-up mechanisms when the competing stimulus representations are perceptually different (Wang, Arteaga, & He, 2013) leaving sensory areas in charge of resolving perceptual conflict when that is not the case (Xu et al., 2016).

### Amygdala and perceptual awareness

In the current study, no evidence of body expression processing outside awareness was found in the amygdala, as its activity was significantly lower than intermediate and clear levels of awareness and was also not significantly different from baseline at PAS1. These findings are therefore in disagreement with previous masking (Morris et al., 1998; Whalen et al., 1998) and binocular rivalry studies (Pasley et al., 2004; Williams et al., 2004) reporting amygdala responses to non-consciously perceived emotional stimuli. Several factors may explain the difference between those studies and the current results.

One explanation may be that these studies often failed to formally evaluate participant’s perception. For example, earlier backward masking studies often determined target discrimination based on subject debriefing after the experiment (Rauch et al., 2000; Sheline et al., 2001; Whalen et al., 1998). In line with this explanation, Pessoa and colleagues (2006) found, after assessing participant’s behavioral performance on a trial-by-trial basis and with SDT measures, differential amygdala activation between fearful and neutral face stimuli only when consciously perceiving the stimuli but not under perceptual unawareness (Pessoa et al., 2006). Similar results have been obtained for other types of threatening stimuli (Hoffmann et al., 2012; Hoffmann et al., 2015). Another possible explanation mentioned earlier could be that previous studies did not account for the different degrees of perceptual awareness, and therefore amygdala activation during reported unawareness could have been confounded with residual awareness.

A complementary explanation relates to the functional heterogeneity of the amygdala (Janak & Tye, 2015; Kyriazi, Headley, & Pare, 2018; LeDoux, 2007; Murray, 2007; Phelps & LeDoux, 2005). In a binocular rivalry study, Lerner and colleagues (2012) found an interesting dissociation in the responses of the different amygdala subregions. While the ventral part of amygdala showed higher responses to unseen fearful faces, dorsal amygdala was more consistently activated by consciously perceived fearful faces (Lerner et al., 2012). Similar results have been shown in the masking study by Etkin and colleagues (2004) . In the current study, amygdala definition was not performed at the sub cluster level, which may partly explain the results.

Furthermore, although amygdala activation has been repeatedly reported for facial expressions, both consciously and non-consciously, evidence for its involvement in body expressions processing is not as consistent, with some studies reporting its involvement (Hadjikhani & de Gelder, 2003; Kret et al., 2011; Poyo Solanas et al., 2020; Sinke et al., 2010; van de Riet et al., 2009) while others failed to show it (Seinfeld et al., 2021; Zhan et al., 2018; Zhan, Goebel, & de Gelder, 2021). Recent proposals have suggested that amygdala’s involvement in emotional processing might be more related to facial properties than to emotion per se. In line with this, two behavioral CFS studies reported that the shorter suppression times observed for fearful faces could be explained by low-level facial features (e.g., the contrast), especially those in the eye region (Gray et al., 2013; Yang et al., 2007). Taken together, the fact that in the current study no amygdala involvement was found during non-conscious processing could relate to the different object category used here.

As mentioned above, amygdala activity increased as the subjective perception of the body stimuli became clearer. The correlation between the magnitude of amygdala activation and the degree of target visibility has previously been reported in the masking study by Pessoa and colleagues (2006) . The fact that stimuli visibility seems to modulate amygdala responses challenges the commonly held view that emotional processing in amygdala is automatic and independent of attention (Dolan & Vuilleumier, 2003; Öhman, 2002). In this regard, the current lack of brain evidence of processing without subjective awareness could be then explained by the fact that body stimuli were presented outside the center of attention (i.e., right or left to the fixation cross). This explanation, however, may not apply in affective blindsight (Morris et al., 2001) or unilateral neglect patients (Vuilleumier et al., 2002). Another explanation already proposed by Pessoa and colleagues (2006) is that the activity of amygdala may itself determine stimulus visibility instead of stimulus awareness being the modulating factor of amygdala’s activity. According to Pessoa and colleagues (2006), this interpretation does not contradict findings in blindsight patients, who may be able to perceive emotional stimuli due to increased amygdala sensitivity to this type of stimuli (Pessoa et al., 2006).

### Pulvinar and perceptual awareness

Although the activity of pulvinar was significantly different from zero in all PAS levels for both emotions, no evidence of emotion or perceptual awareness modulations were found in this area. This contradicts the findings by Padmala et al. (2010) showing a correlation between pulvinar responses and stimulus detection, especially for affectively conditioned stimuli (Padmala, Lim, & Pessoa, 2010). Previous research in human and non-human primates has shown that this area is constituted by retinotopic maps of the contralateral visual fields. The fact that the presentation of the target stimuli was performed to either the right or left visual field may have ‘diluted’ any possible perceptual awareness or affective modulations. Another explanation could be that in the current experiment we did not differentiate between false alarms and hit trials. In a face detection task study, Koizumi and colleagues (2019) reported higher pulvinar activity for false alarm trials than hit trials independent of emotion (fearful or happy faces) (Koizumi et al., 2019). Also, in that study no significant differences between fearful and happy hit trials were reported. Taken together, those results point to a general role of pulvinar in signaling false percepts, which was not taken into account in the current study.

### Affective processing and perceptual awareness

As mentioned earlier, participants’ ability to discriminate fearful from neutral bodies increased linearly with increased perceptual awareness. Moreover, sensitivity values differed from chance performance at all PAS levels except for ‘no experience’. While these results indicate that emotional discrimination may not be observed during perceptual unawareness in neurologically intact participants (Rajananda et al., 2020), it is crucial to note that our results, like any others, are narrowly linked to the methods used to create (un)awareness. CFS is among the best methods to control for visual awareness, yet we do not fully understand how it interferers with the normal processing flow of feedforward and feedback projections in the visual system, including cortical as well as subcortical information streams.

At the physiological level, fearful bodies triggered a slower heart rate compared to neutral ones for ‘almost clear’ stimulus experience (**Figure 5**). This is in line with freezing behavior, characterized by a deceleration of heart rate when confronted with a threat (Roelofs, 2017). Heart rate modulations have shown to depend on several factors, such as the ambiguity of the threat. During the almost clear experience of the stimulus, stimulus visibility may have been enough to recognize a body (or parts of it) but not sufficient to be completely certain of the emotional expression. This perceptual ambiguity may explain why freezing behavior was most pronounced for fearful body expressions when participants reported an ‘almost clear’ experience of the stimulus.

At the brain level, the activity in the areas showing an effect of perceptual awareness was not influenced by emotion. Previous studies showing emotion influences in fronto-parietal areas used facial expressions (Whalen et al., 1998), reported an area location that did not correspond to the ones in the current study (IFC, IPS) (Amting, Greening, & Mitchell, 2010; Vuilleumier et al., 2002) or included large parts of the fronto-parietal cortices (Kiss & Eimer, 2008), which may explain the current results. Future studies using other emotional stimuli than facial expressions are needed to investigate whether the resolution of perceptual transitions in fronto-parietal cortices occurs independently of the emotional content. Furthermore, no emotional modulation was found in high-level visual cortices in the current study. The involvement of EBA and FG in body expression perception is still inconclusive (van de Riet et al., 2009), with some studies supporting their role (Hadjikhani & de Gelder, 2003; Kret et al., 2011; Pichon, de Gelder, & Grèzes, 2009) while others ascribe it to attention and arousal confounding factors (Downing & Peelen, 2011).

In the case of the amygdala, higher activity was found for neutral body expressions than fearful ones, in disagreement with previous literature linking this region with the processing of threatening signals (Adolphs, 1999). Although participants were able to correctly distinguish neutral bodies from fearful ones when reporting higher stimulus visibility (during almost clear and clear trials), it could be that the action represented by the neutral body posture (opening door) was more ambiguous than the fearful expression, which may explain the results. This interpretation agrees with previous studies suggesting this area as key in the signaling and resolution of ambiguity (Adolphs, 2002; de Gelder et al., 2014; Hortensius et al., 2017; Poyo Solanas et al., 2018; Whalen et al., 1998).

## Conclusions

The current study provides behavioral, physiological and neuroscientific evidence of the processing of threatening body expressions at 7T using a finer measure of perceptual awareness in combination with signal detection theory measures. The use of the PAS scale revealed intermediate states of perceptual awareness and allowed to differentiate non-conscious processes from partial awareness states. In particular, this study shows a gradual relationship between recognition sensitivity and perceptual awareness as well as no evidence of perceptual discrimination during perceptual unawareness. In addition, it shows that there are both gradual and dichotomous relationships between perceptual awareness and brain activity, possibly reflecting the stimuli or feature processing stage or the different function of each area.

## Materials and Methods

### Participants

Fifty-one healthy volunteers were recruited in this study. However, only seventeen healthy volunteers (mean age = 20.69 years; age range = 19-29 years; 11 female; all right-handed) met the required criteria (see *Experimental design and procedure* section) and participated in the fMRI experiment. Participants had normal or corrected-to-normal vision and a medical history without any psychiatric or neurological disorders. The experiment was approved by the Ethical Committee at Maastricht University and was performed in accordance with the Declaration of Helsinki. Participants provided informed written consent before the start of the experiment and received vouchers or credit points after their participation. In addition, participants remained unaware of the aim of the study until the completion of the experiment and were unfamiliar to the CFS paradigm.

### Experimental design and procedure

Each participant took part in two scan sessions performed on separate days and in randomized order. In one of the sessions, six functional runs of the main CFS experiment were acquired as well as the anatomical data of the participant (∼2h 15min). In the other session, resting state data, the data of a body-area localizer and a population receptive field localizer for motion-sensitive early- and mid-level visual cortex were acquired (∼1.5h) see *Supplementary Information* for more information on this session). Before the participation in the scanning sessions, participants underwent a short behavioral experiment (∼30min) on a separate day to ensure their eligibility for the CFS experiment. Participants showing unstable merging of the stimuli and/or strong suppression were excluded from taking part in the fMRI sessions. The behavioral session also served to determine eye-dominance under CFS (see *Supplementary Information* for more information on the behavioral session).

In the main CFS experiment, the participants’ non-dominant eye was presented with a static body posture while a colorful Mondrian mask flickering at 10 Hz was presented to the dominant eye. Dichotomous presentation was accomplished using a cardboard panel and a pair of prism glasses. The cardboard was placed between the mirror attached to the head coil and the screen, dividing it into two halves and ensuring that each eye only perceived half of the screen. The prism glasses (diopter = 6) bent the light in a way that the ipsilateral image was shifted back to the center of each eye (as described in Schurger, 2009). Both the body stimuli and the colorful mask were displayed on a grey background (RGB value = 128, 128, 128) within a black rectangular frame (frame thickness=10 pixels, frame size 318×352 pixels, 5.08°x5.62° visual angle) that had a fixation cross at its center, respectively, which facilitated the merging of the two images (see **Figure 1**).

The body stimuli were selected from a large validated stimulus set of still whole-body images (Stienen & de Gelder, 2011) and consisted of eight actor identities (half females) that portrayed either a fearful or a neutral (opening door) body expression (318×182 pixels, 5.08°x2.91° visual angle), with the facial information removed. The body postures were presented either to the right or left side of the fixation cross in a randomized order. The colorful Mondrian mask consisted of 600 unique patterns flashing randomly at 10Hz, which were comprised of overlapping small rectangles covering the entire rectangular frame (see **Figure 1**).

Once the participant reported stable perception of a single rectangle, the experimental run started with a twelve-second fixation period. A change in the fixation cross color from black to white indicated the start of each trial and remained white for the whole trial duration. Each trial started with a one-second white fixation period, followed by a two-second CFS presentation consisting of a gradual increase of the body stimulus contrast over one second, followed by the ramp down of the contrast back to 0% within 0.5s, and a 0.5s blank period. The gradual increase of the stimulus contrast was performed to decrease the likelihood of the body stimulus escaping suppression. The contrast of the colorful Mondrian mask was constant throughout the two-second CFS presentation within each trial. However, both the contrast of the body stimuli and the Mondrian mask were determined for each trial using a staircase procedure with 10 steps (body stimuli: 5%, 14%, 23%, 32%, 41%, 50%, 50%, 50%, 50%, 50%; noise: 100%, 100%, 100%, 100%, 100%, 82%, 64%, 46%, 28%, 10%) that depended on the participant’s visual experience of the body stimulus in the previous trial. If participants reported not seeing anything in the colorful noise, the maximum contrast of the body stimuli increased one step while the contrast of the mask decreased, also by one step. Each run started at step 5 (i.e., 41% contrast for body stimulus and 100% for the noise). This staircase procedure was intended to balance the number of trials per perceptual awareness condition.

The two-second CFS period was followed by a jittered fixation period (4-6-8s) after which participants were required to make two responses. The first response required participants to categorize the body stimulus in a two-alternative forced-choice manner (fearful vs. neutral) by pressing one of two buttons. The assignment of the two buttons was randomized. Subsequently, participants had to indicate their visual experience of the stimulus according to the perceptual awareness scale by pressing one out of four buttons: ‘no experience’ (PAS1), ‘brief glimpse’ (PAS2), ‘almost clear experience’ (PAS3) and ‘clear experience’ (PAS4). The button assignment was kept constant for this task to facilitate a quick response. Both responses were required even when participants reported not seeing anything in the noise. In those cases, participants were instructed to guess the emotional expression of the stimulus. Both answers had to be given within a 1.5-second window each, and always with the right hand. Participants were informed about the short response window during the preceding behavioral session, where two practice runs were administered (see *Supplementary Information*). In addition, participants were instructed to keep as still as possible throughout the experiment, to always fixate on the cross and not to blink within the two-second CFS period. Each response period was followed by a jittered inter-trial-interval (4-6-8s), resulting in an average trial duration of 18s and an average run duration of 10min approximately. Each run was comprised of 32 trials, 16 per emotional condition, with two repetitions for each of the eight body stimulus identities. Therefore, a total of 192 trials were obtained, 96 for each emotional category. One participant only performed four runs due to delays in the scanning. Two participants performed seven runs instead of six. One run of three participants and two runs of another were discarded due to excessive motion.

The experiment was presented in MATLAB R2012a (MathWorks, Natick, MA, USA) using Psychotoolbox 3.0.11 (Brainard & Vision, 1997; Pelli & Vision, 1997). The stimuli were back-projected on a translucent screen situated at the end of the scanner bore, behind participants’ heads (Panasonic PT-EZ570; Newark, NJ, USA; screen size = 30 x 18 cm, screen resolution = 1920 x 1200 pixels, refresh rate = 60 Hz, visual angle = 17.23° x 10.38°). Participants viewed the screen through a tilted mirror attached to the head coil. The distance between the mirror and the screen was ∼99 cm. Participant responses were recorded using an MR-compatible button box (Current Designs, 30 8-button response device, HHSC-2 × 4-C; Philadelphia, USA).

### Behavioral data analysis

Behavioral data were analyzed with SPSS (version 22.0; IBM Corp., Armonk, N.Y., USA) and custom code in MATLAB R2020a (MathWorks, Natick, MA, USA). First, trials without a response for one or both tasks were excluded from further analyses. Trials in which reaction times deviated more than 3.5 times the standard deviation from the mean (within run and subject) were also removed. In total, 136 out of 3239 trials (4.2%) were excluded from further analyses.

Participants’ responses in the two-alternative forced-choice task were counted as hits (H), misses (M), correct rejections (CR) and false alarms (FA) according to Signal Detection Theory (Green & Swets, 1966; Tanner & Swets, 1954). Hits refer to trials in which fearful bodies were correctly categorized, while misses to those trials in which participants incorrectly categorized fearful bodies as neutral. Correct rejections indicate trials where neutral bodies were correctly categorized whereas false alarms to the trials where neutral stimuli were incorrectly categorized as fearful ones (Candidi, Stienen, Aglioti, & de Gelder, 2011; Snodgrass & Corwin, 1988; Tamietto, Geminiani, Genero, & de Gelder, 2007).

To further understand participants’ responses, the perceptual sensitivity (d’) and the response criterion or bias (c) were calculated for each PAS level. Sensitivity is commonly calculated by subtracting the z-transformed false alarm rates from the z-transformed hits (**Equation 3**), and therefore reflects the distance between the target (fearful body) and noise (neutral body) distribution means, in standard deviation units. Here, a modified form of hit (H’) and false alarm (FA’) rates was used to account for ceiling effects, as proposed by Snodgrass & Corwin (1988) (**Equation 1 & 2**). Higher sensitivity values indicate higher discriminability of fearful bodies from neutral ones. A value of zero indicates inability to distinguish fearful body expressions from neutral ones. Independent from sensitivity, criterion bias was calculated by multiplying the sum of the z-transformed hit and false alarm by 0.5 (**Equation 4**) (Macmillan, 1993; Snodgrass & Corwin, 1988; Tamietto et al., 2007). It reflects the distance between the neutral point (where responses are not biased towards fearful bodies nor neutral ones) and the response criterion, in standard deviation units. Negative response criterion values indicate a bias in reporting the presence of a fearful body over a neutral one (liberal criterion), while positive values show the opposite response pattern (conservative criterion).

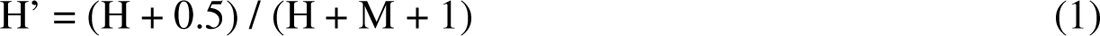

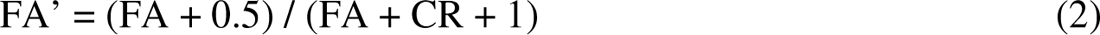

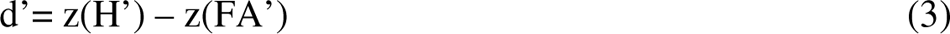

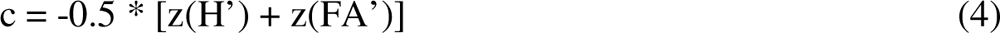

Average sensitivity and criterion bias values were calculated for each perceptual awareness rating and participant. Subsequently, sensitivity and criterion bias values were analyzed, respectively, using a linear mixed model procedure with the within-subject factor Perceptual Awareness (four levels: PAS1, PAS2, PAS3 and PAS4) and the Toeplitz covariance matrix for repeated measures based on Akaike information criterion (AIC) values (Akaike, 1974). The weighted least squares method was used to account for violations of homoscedasticity in the analysis of sensitivity values. Sensitivity and criterion bias values were also compared to chance level (i.e., against zero) using a one sample t-test per perceptual awareness level.

Next, to investigate whether perceptual awareness is a gradual or a dichotomous phenomenon, we fitted two linear mixed models to the sensitivity data with different predictor definitions. In the gradual model, the predictors modelled a linear relationship between sensitivity and the PAS levels. In the dichotomous model, the predictor for the PAS1 level was set to zero while the rest of the PAS levels were set to 1, describing an ‘all-or-none’ relationship between recognition sensitivity and perceptual awareness. These two models were performed independently for each participant. To select the model that best represented the recognition sensitivity pattern across PAS levels at the group level, the values corresponding to the Bayesian information criterion (BIC) (Stone, 1979) resulting from each model fitting were analyzed with a paired-sample t-test (gradual vs. dichotomous). The model with the significantly lower BIC value (which indicates better fit) was selected as the final model.

Lastly, the reaction times (RTs) of the emotional recognition task were analyzed using a linear mixed model with SDT (four levels: H, M, FA, CR) as the within-subject factor. The RTs of the perceptual awareness task were also analyzed with a linear mixed model with Perceptual Awareness (four levels: PAS1, PAS2, PAS3 and PAS4) as the within-subject factor. Both models used the Toeplitz covariance matrix for repeated measures.

### (f)MRI data acquisition

(f)MRI data were acquired with a 1-transmitter/32-receiver head coil (NovaMedical Inc.; Wilmington, MA, USA) in a 7 Tesla Magnetom whole-body scanner (Siemens Medical Systems, Erlangen, Germany) located at the Maastricht Brain Imaging Centre (MBIC), the Netherlands. Functional images were obtained using a 2D gradient echo (GE) echo-planar imaging (EPI) sequence (voxel size = 1.2 mm isotropic, no gap, repetition time (TR) = 2000 ms, echo time (TE) = 21 ms, flip angle (FA) = 75°, in-plane field of view (FoV) = 172.8 x 172.8 mm^2^, matrix size = 144 x 144, number of slices per volume = 70, multiband acceleration factor = 70, multiband acceleration factor = 2, iPAT=3, phase encoding direction = anterior to posterior, bandwidth=1488 Hz/Px, echo spacing=0.78 ms, number of volumes = 300 (main experimental runs), 440 (body-areas localizer), 315 (pRF mapping), 330 (resting state)). The slice positioning of the functional images was performed in a way to include the occipital, parietal and frontal lobes as well as the amygdala, thus ensuring a good coverage of important areas in body perception. However, limited coverage was obtained for the superior part of the motor cortex, anterior temporal lobe and orbito-frontal cortex. For distortion correction of the functional images, a short run (5 volumes) was acquired before each experimental run with the same parameters specified above but with opposite phase encoding direction (posterior-to-anterior). Anatomical images were acquired for each participant using a MP2RAGE sequence (voxel size = 0.65 mm isotropic, FoV = 207 x 207 mm^2^, 2.51 ms, Inversion Time (IT) 1 = 900ms, IT2 = 2750 ms, FA1L 5°, FA2 = 3°, iPAT = 2, bandwidth=250 Hz/Px, echo spacing = 7 ms). Dielectric pads covering the occipital and temporal lobes were used for all participants.

### (f)MRI data preprocessing

The pre-processing and analysis of the (f)MRI data were performed in BrainVoyager (v22.0; Brain Innovation B.V.) as well as with custom codes in MATLAB (vR2020a; The MathWorks Inc.; Natick, MA, USA). First, functional images underwent top-up distortion correction with the COPE (Correction based on Opposite Phase Encoding) plugin (v1.1.1) in BrainVoyager (Heinrich, Papież, Schnabel, & Handels, 2014) based on the voxel displacement between the first volume of the functional run and that of the distortion correction run. Subsequently, slice scan time correction was applied to the functional runs using sinc interpolation. Functional images then underwent 3D rigid motion correction with respect to the first volume of each functional run (trilinear/sinc interpolation). Linear trend removal and high-pass temporal filtering were employed to exclude low-frequency drifts using a general linear model (GLM) Fourier basis set with 2 cycles per time course. In order to reduce the B1 bias field, the anatomical data were background-noise corrected by dividing the UNI image by the T1w image and then masking the resulting ratio image by the INV2 image. In addition, the structural data were corrected for intensity inhomogeneities and upsampled to 0.6 mm isotropic resolution (sinc interpolation, framing cube = 384) to best match the resolution of the functional data. After these steps, each pre-processed functional run was aligned to the first run of the main experiment and normalized to Talairach space. The resulting co-registered images were then spatially smoothed with a Gaussian kernel of a full-width half-maximum of 3mm. A group-averaged anatomical image was created by averaging the Talairach-normalized anatomical data across participants. All analyses were performed in volume space.

To facilitate the visualization of group results, a cortex-based alignment (CBA) procedure was carried out. First, the T1-weighted anatomical data of each participant were downsampled to 0.7mm isotropic (for better software results) and subsequently underwent a DNN-based segmentation procedure in BrainVoyager (strides value slow: 32×32×32). This approach classified each anatomical voxel into eight possible tissue types, including white matter, grey matter, cerebrospinal fluid, blood vessels, ventricles, subcortical structures, sagittal sinus and background. With this information, all the individual anatomical UNI datasets were then segmented at the grey– white matter boundary, upsampled to 0.6mm isotropic and normalized to Talairach space. After this step, manual corrections were performed when necessary, on a slice-by-slice basis. The cortical surfaces were then reconstructed, inflated, smoothed, and mapped onto a high-resolution standard sphere, separately for each hemisphere (vertices = 163842). A dynamic group averaging approach based on individual curvature information was used to align participants’ reconstructed cortical surfaces. After alignment, an averaged folded cortical mesh (*n*L L each hemisphere.

### Physiological data acquisition, preprocessing and noise correction of fMRI data

Cardiac and respiratory measures were acquired using an oximeter (50Hz) and a pneumatic compression belt (50Hz) to control for physiological fluctuation effects on the BOLD response (Birn, Smith, Jones, & Bandettini, 2008; Chang, Cunningham, & Glover, 2009; Glover, Li, & Ress, 2000; Shmueli et al., 2007). Low-frequency drifts were removed from the raw cardiac (bandpass, 0.5-8Hz) and respiratory (lowpass, 2Hz) data and signal peaks were identified (Elgendi, Norton, Brearley, Abbott, & Schuurmans, 2013). Physiological noise correction was performed using a modification of the conventional Retrospective Image Correction (RETROICOR) procedure (Glover et al., 2000; Harvey et al., 2008; Hu, Le, Parrish, & Erhard, 1995; Hutton et al., 2011). In the current procedure, a cardiac and a respiratory phase were assigned to each functional image using third-order cardiac and fourth-order respiratory harmonics. The respiratory phase not only considered the respiratory timing but also the depth of the breathing (Glover et al., 2000). In addition, a cardiorespiratory interaction (first order) term was defined (Harvey et al., 2008). A total of 20 regressors were created, including 6 cardiac phase regressors, 8 respiratory phase regressors, 4 cardiorespiratory interaction regressors, as well as a filtered heart rate and respiratory rate regressor.

### (f)MRI data analysis

#### Definition of regions of interest sensitive to perceptual awareness

A whole-brain fixed-effects GLM was performed for each subject, individually, with the 3mm-smoothed percent-signal normalized functional data. The GLM included as predictors of interest four predictors corresponding to the perceptual awareness levels (i.e., PAS1-4), a parametric predictor of the mask contrast, a predictor for ‘no response’ trials and a predictor for each of the two response windows. These predictors were convolved with a two-gamma hemodynamic response function. In addition, six motion predictors and 20 physiological predictors (see previous section) were included in the design matrix as nuisance predictors. For each subject, a beta map for each perceptual awareness condition was obtained and entered into a group repeated-measures ANOVA in BrainVoyager. One subject was excluded from this analysis due to a missing condition (PAS4). The resulting t-map showing the main effect of PAS at the group level was corrected for multiple comparisons using a cluster-threshold procedure based on Monte-Carlo simulations (initial p-value = .001, alpha level = .05). This map was then used to define regions of interest (ROI) in each subject for subsequent analyses.

#### Anatomical definition of amygdala and pulvinar

In addition to the PAS-sensitive defined ROIs, the pulvinar and the amygdala were defined in each subject using the Chakravarty Atlas (Chakravarty, Bertrand, Hodge, Sadikot, & Collins, 2006) given their involvement in non-conscious processing (Tamietto & de Gelder, 2010). The definition of pulvinar covered all different subnuclei of both the left and right hemisphere. Amygdala definition also covered all subnuclei of both the left and right hemisphere and was manually modified according to individual anatomy when necessary.

#### Analysis of the defined ROI data

A fixed-effects GLM similar to the one described above was performed for each subject with the 3mm-smoothed functional data. However, the predictors corresponding to the perceptual awareness levels were now separated according to the emotional category of the stimulus (i.e., N1, N2, N3, N4, F1, F2, F3, F4; N = neutral; F = fear). For each subject, the beta values corresponding to these eight main conditions were extracted from each ROI (functionally and anatomically defined) and entered into a linear mixed model analysis in SPSS. This analysis was performed for each ROI separately and included two within-subject factors: Emotion (two levels: neutral, fear) and Perceptual Awareness (four levels: PAS1, PAS2, PAS3, PAS4). All analyses used the unstructured covariance matrix for repeated measures. Multiple comparisons were corrected within each ROI with the Sidak-method and across ROIs with the Benjamini-Hochberg false discovery rate (BHFDR) method. To examine whether ROI activity was consistently above or below baseline, a one-sample *t*-test against 0 was performed for each of the eight experimental conditions within each ROI (FDR correction at *q* < .05).

As with the sensitivity data, we conducted further analyses to investigate whether brain activity showed a gradual or a dichotomous relationship to perceptual awareness. This resulted in the fitting of four linear mixed models (two Models: Gradual and Dichotomous; two Emotions: Neutral and Fear) into the data of each ROI and subject, respectively. The resulting BIC values from each model fitting were entered into a repeated measures ANOVA with within-subjects factor Model (two levels: gradual and dichotomous) and Emotion (two levels: neutral and fear). In the cases where there was a significant effect of Model but not a significant Model*Emotion interaction, a paired t-test was performed between the coefficient estimates of the neutral and fearful models to assess how different the model slopes and intercepts were across emotions. In the cases where there was a significant Model*Emotion interaction, a different model was selected for each emotion. When no significant effect of Emotion and Model were found, as well as no significant interaction, two model fittings were performed (Gradual and Dichotomous) after averaging the ROI data across emotions. Subsequently, a paired-sample t-test was performed with the resulting BIC values from each model fitting. The BHFDR method was used to correct for multiple comparisons across ROIs and the Sidak-method to correct them within each ROI.

### Cardiac data analysis

For each trial, the systolic peaks corresponding to the two-second CFS period were identified. Peaks beyond a biologically feasible range were rejected (i.e., beats per minute < 35 or > 180) as well as outliers that were 2.5 standard deviations from the mean. Subsequently, the mean heart rate (beeps per minute) was obtained by averaging the time differences between consecutive peaks. These average estimates were baseline-corrected with respect to the mean heart rate corresponding to the one second preceding each CFS period. The resulting values were entered into a linear mixed model procedure with within-subject factors Emotion (two levels: anger and fear) and Perceptual Awareness (four levels: PAS1, PAS2, PAS3 and PAS4). The compound symmetry covariance matrix for repeated measures was used. Two outliers (single data points within the whole sample) were removed based on their standardized residuals resulting a model with significantly better fit.

## Acknowledgments

This work was supported by the European Research Council (ERC) FP7-IDEAS-ERC (Grant agreement number 295673; Emobodies), by the ERC Synergy grant (Grant agreement 856495; Relevance), by the Future and Emerging Technologies (FET) Proactive Program H2020-EU.1.2.2 (Grant agreement 824160; EnTimeMent) and by the Industrial Leadership Program H2020-EU.1.2.2 (Grant agreement 825079; MindSpaces). We would like to thank J. Eck for the help regarding the physiological noise correction of fMRI data, to M. J. Vaessen for the help during the piloting of this study and to V. Smekal for comments on an earlier draft of the introduction section.

## Competing interests

The authors declare no competing interests.

## Author contributions

Conceptualization: M.P.S., M.Z. and B.d.G.; Methodology and Software: M.P.S. and M.Z.; Investigation & Formal Analysis: M.P.S.; Writing – original draft: M.P.S.; Writing – review & editing: M.P.S., M.Z. and B.d.G; Visualization: M.P.S.; Funding Acquisition: B.d.G.

## Notes

### Competing Interest Statement

The authors have declared no competing interest.

