## Supplementary for "Perceptual awareness is gradual in temporal and dichotomous in fronto-parietal cortices"

### Supplementary Information

#### Supplementary Materials

##### *Behavioral session*

Before the participation in the scanning sessions, participants underwent a short behavioral experiment (~30min) in a separate day to ensure their eligibility for the experiment. First, an eye-dominance test was administered to determine the participant's dominant eye. Two practice runs were subsequently administered if the participant showed stable merging during the eye-dominance test as well as not a strong suppression. Participants that did not meet these criteria were excluded from further participation in the study and their data were discarded. The same assessment procedure was performed after the practice runs. Therefore, only participants that met these criteria for both the eye-dominance and practice runs were contacted for the fMRI sessions.

*Eye dominance test*. The dichotomous presentation was achieved in the same manner as in the main experiment using a cardboard and prism glasses, although outside the scanner. For this test, ten different neutral faces (half male) belonging to the Radboud Face Database (Langner et al., 2010) were selected. The face stimuli (318x212 pixels, 5.08°x3.39° visual angle) were presented to one eye in the center of a rectangular frame (318x212 pixels, 5.08°x3.39° visual angle, 10 pixels wide) while a dynamic colorful mask pattern (318x212 pixels, 5.08°x3.39° visual angle) flashing at 10Hz, and covering the other entire rectangular frame, was shown to the other eye. Each trial consisted of a gradual ramping up of the face stimulus contrast from 0% to full contrast over one second, which was maintained for another second and then followed by the diminishment of the stimuli contrast to 0% over 0.5s and a 0.5s blank period. During these three seconds, the contrast of the dynamic colorful mask remained constant. Next, a fixation dot appeared in the screen indicating participants to report whether they saw or did not see a face by pressing one out of two keys ('J' for seen, 'K' for unseen). Each stimulus was randomly presented three times to each eye, giving a total of 60 trials. Eye dominance was defined as the eye that perceived the highest amount of seen trials while it was assigned randomly in the cases where the amount of seen trials was equal between both eyes.

*Practice runs.* Two practice runs were administered with an identical experimental design to the main task of the fMRI session. The only difference was a shorter inter-trial interval (ITI) and a shorter interval between the two-second CFS period and response periods. The aim of these practice runs was to familiarize participants with the perceptual awareness scale as well as to train them to respond as accurate and fast as possible given the short response window (~1.5 seconds) for each of the tasks.

***Second fMRI session***

Each participant took part in two scan sessions performed in separate days and in randomized order. The session corresponding to the main experimental task is described in the main text. In the other session, resting state data, the data of a body-area localizer and a population receptive field localizer for motion-sensitive early- and mid-level visual cortex were acquired (~1.5h). The body-area localizer experiment was presented in MATLAB vR2012a (MathWorks, Natick, MA, USA) using Psychtoolbox v3.0.11 (Brainard & Vision, 1997; Pelli & Vision, 1997) while the MT localizer was presented in PsychPy (v1.90.0) (Peirce, 2007, 2009). The stimuli were back-projected on a translucent screen situated at the end of the scanner bore, behind participants' heads

(Panasonic PT-EZ570; Newark, NJ, USA; scree size = 30 x 18 cm, screen resolution = 1920 x 1200 pixels, refresh rate = 60 Hz, visual angle = 17.23° x 10.38°). Participants viewed the screen through a tilted mirror attached to the head coil. The distance between the mirror and the screen was of ~99 cm. Participant responses were recorded using an MR-compatible button box (Current Designs, 30 8-button response device, HHSC-2 × 4-C; Philadelphia, USA). The data resulting from these runs was not analyzed in the current study.

*Body-area localizer.* This localizer employed a block design and included still grey-scale images of bodies, faces, houses, tools and words. Body images consisted of 18 actor identities (nine female), with the facial information removed, portraying a neutral expression (de Gelder & Van den Stock, 2011). Facial images depicting a neutral expression (18 identities, nine female) were taken from the Karolinska Directed Emotional Faces (KDEF) database (Lundqvist, Flykt, & Öhman, 1998). Tool (18 hand-held tools) and house (18 house facades) stimuli were obtained from the internet. Word stimuli consisted of high-frequency 4-6 letter words in Arial font. All images were presented on a grey background (RGB value =157, 157, 157) and spanned 3.5 x 3.5° of visual angle. The experiment started with a 12-second fixation period. Each block had a duration of 12 seconds in which 12 images of the same category were presented in randomized order for 800ms with an inter-stimulus interval of 200ms. Each block was separated by a 12-second fixation period. Each category was presented seven times under passive-viewing conditions in a pseudo-randomized order. The total duration of this localizer was approximately 14 minutes.

*pRF mapping.* A population receptive field localizer for motion-sensitive early- and mid-level visual cortex was used (https://github.com/MSchnei/pyprf_motion). Mapping stimuli consisted of bars or wedge apertures containing a 'ripple' pattern used in previous studies (Alvarez, de Haas, Clark, Rees, & Schwarzkopf, 2015; Schwarzkopf, Anderson, de Haas, White, & Rees, 2014; van

Dijk, de Haas, Moutsiana, & Schwarzkopf, 2016). Bars were oriented either vertically or horizontally and subtended a width of 3 degrees of visual angle while the wedges subtended 45°. The experiment was constituted of four randomized experimental blocks (one vertical bar, one horizontal bar and two wedges blocks), where carriers of the same type were randomly presented two times at 16 different locations for 0.5s (ITI = 1.2s) within a circular grid (grid radius = 12°). Experimental blocks were separated by rest periods (Repetition Time (TR) = 14) when only the grid was shown. At the center of the grid there was a fixation dot constituted by a red circle (radius = 0.125°) surrounded by a yellow annulus (radius = 0.19°). Participants were asked to fixate on the dot throughout the entire run as well as to indicate a color change of the fixation dot by pressing a button. The colour change was always from red to yellow for 0.3s. The total duration of this run was approximately 10 minutes.

*Resting state.* A resting state run was acquired during with a duration of 11 minutes. During this run, participants were instructed to fixate at a white cross located at the center of the screen (background RGB = 0, 0, 0) and not to think of anything specific.

#### Supplementary Results

**Table S1. Mean reaction times and standard errors of the emotional recognition task (RT-emo) and the Perceptual Awareness rating (RT-pas)**

| | **HIT** | **MISS** | **FA** | **CR** |
|---|---|---|---|---|
| **RT-emo** | 0.77 ± 0.03 | 0.75 ± 0.03 | 0.75 ± 0.03 | 0.77 ± 0.03 |
| | **PAS1** | **PAS2** | **PAS3** | **PAS4** |
| **RT-pas** | 0.48 ± 0.04 | 0.56 ± 0.04 | 0.55 ± .04 | 0.58 ± 0.04 |

*Note*: mean reaction time values are estimated marginal means. *Abbreviations*: CR: correct rejection; FA: false alarm; PAS: perceptual awareness scale; RT-emo: reaction times of the emotional categorization task; RT-pas: reaction times of the visual awareness rating.

**Table S2. Mean percentage of responses and standard error separated by Signal Detection Theory measures and Perceptual Awareness ratings**

| | **PAS1** | **PAS2** | **PAS3** | **PAS4** |
|---|---|---|---|---|
| **HIT** | 0.05 ± 0.02 | 0.08 ± 0.02 | 0.09 ± 0.02 | 0.06 ± 0.02 |
| **MISS** | 0.14 ± 0.04 | 0.06 ± 0.01 | 0.02 ± 0.02 | 0.00 ± 0.00 |
| **FA** | 0.05 ± 0.02 | 0.04 ± 0.02 | 0.02 ± 0.01 | 0.01 ± 0.00 |
| **CR** | 0.13 ± 0.04 | 0.12 ± 0.02 | 0.08 ± 0.02 | 0.05 ± 0.02 |

*Abbreviations*: CR: correct rejection; FA: false alarm; PAS: perceptual awareness scale.

**Table S3. Information details of the clusters resulting from the group ANOVA with within-subject factor Perceptual Awareness Scale (four levels: PAS1-4)** (cluster size corrected with Monte-Carlo simulation, alpha level = 0.05, initial p = 0.001, numbers of iterations = 5000).

| | H | Average Talairach coordinates | | | | | | Cluster size ($mm^3$) | Avg. t-value | Avg. p-value |
|---|---|---|---|---|---|---|---|---|---|---|
| | | X | X (SD) | Y | Y (SD) | Z | Z (SD) | | | |
| **ITG** | R | 98 | 2.99 | -72 | 3.41 | -18 | 3.11 | 976 | 7.85 | .000 |
| **LOTC** | R | 83 | 7.54 | -103 | 11.72 | 0 | 11.24 | 25277 | 8.44 | .000 |
| **FG** | R | 70 | 6.37 | -75 | 10.60 | -24 | 5.59 | 21334 | 10.31 | .000 |
| **AMYG** | R | 36 | 3.85 | -10 | 2.20 | -14 | 2.36 | 565 | 7.63 | .000 |
| **Precuneus** | L | -3 | 7.51 | -112 | 5.68 | 68 | 3.13 | 2637 | 7.77 | .000 |
| **Fourth OG** | L | -11 | 3.14 | -148 | 2.34 | -20 | 1.56 | 805 | 8.26 | .000 |
| **SOG** | L | -17 | 2.63 | -158 | 2.12 | -6 | 2.85 | 762 | 7.65 | .000 |
| **Fourth OG** | L | -33 | 4.97 | -132 | 7.58 | -20 | 3.15 | 6072 | 8.95 | .000 |
| **aIPS** | L | -70 | 6.38 | -66 | 7.15 | 73 | 3.93 | 6283 | 7.95 | .000 |
| **mIPS** | L | -52 | 4.98 | -94 | 9.70 | 65 | 6.53 | 6316 | 7.52 | .000 |
| **pIPS** | L | -48 | 2.31 | -121 | 4.96 | 50 | 3.79 | 1277 | 7.59 | .000 |
| **pSTS** | L | -82 | 9.88 | -111 | 12.08 | -1 | 11.90 | 28866 | 8.49 | .000 |
| **LOTC** | L | -79 | 9.15 | -85 | 3.57 | 17 | 7.93 | 6444 | 7.58 | .000 |
| **PMv** | L | -76 | 7.74 | 19 | 8.29 | 53 | 5.64 | 9395 | 8.33 | .000 |
| **FG** | L | -69 | 4.90 | -78 | 13.88 | -24 | 5.29 | 18894 | 9.46 | .000 |
| **ITG** | L | -92 | 4.26 | -68 | 4.44 | -19 | 2.95 | 3036 | 8.13 | .000 |
| **STG** | L | -98 | 2.10 | -25 | 1.43 | 14 | 1.57 | 347 | 8.13 | .000 |

*Abbreviations*: aIPS: anterior intraparietal sulcus; AMYG: amygdala; Avg.: average; F: fear; FG: fusiform gyrus; H = hemisphere; IFC: inferior frontal cortex; ITG: inferior temporal gyrus; LOTC: lateral occipito-temporal cortex; mIPS: medial intraparietal sulcus; N: neutral; N.A.: not applicable; OG: occipital gyrus; PAS: perceptual awareness scale; pIPS: posterior intraparietal sulcus; pSTS: posterior superior temporal sulcus; SD = standard deviation; STG: superior temporal gyrus.

**Table S4. Results of the linear mixed model analysis with within-subject factors Emotion and PAS, separately for each ROI.**

| | PAS effect | Emotion effect | Interaction effect | PAS pairwise comparisons | | | | | |
|---|---|---|---|---|---|---|---|---|---|
| | | | | 4>1 | 4>2 | 4>3 | 3>2 | 3>1 | 2>1 |
| **ITG [r]** | F(3,12.57) = 16.11, p < .001, pcorr < .001 | F(1,12.25) = 1.23, p = .288, pcorr = .828 | F(3,10.89) = 1.36, p = .307, pcorr = .651 | *** | * | | * | ** | |
| **LOTC [r]** | F(3,13.71) = 21.09, p < .001, pcorr < .001 | F(1,10.68) = 2.06, p = .179, pcorr = .762 | F(3,11.07) = 0.78, p = .529, pcorr = .881 | *** | * | | ** | *** | * |
| **FG [r]** | F(3,14.91) = 13.88, p < .001, pcorr < .001 | F(1,11.73) = 0.02, p = .879, pcorr = .879 | F(3,12.38) = 0.59, p = .634, pcorr = .881 | ** | | | * | *** | * |
| **AMYG [r]** | F(3,16.01) = 8.51, p = .001, pcorr = .002 | F(1,14.63) = 0.16, p = .691, pcorr = .828 | F(3,15.49) = 0.52, p = .674, pcorr = .881 | ** | † | | | | * |
| **Precuneus [l]** | F(3,16.03) = 13.05, p < .001, pcorr < .001 | F(1,15.37) = 0.70, p = .415, pcorr = .828 | F(3,15.69) = 0.68, p = .578, pcorr = .881 | ** | | | | *** | † |
| **Fourth OG [l]** | F(3,14.19) = 6.13, p = .007, pcorr = .007 | F(1,14.80) = 0.03, p = .873, pcorr = .879 | F(3,13.13) = 0.22, p = .882, pcorr = .937 | ** | | | | | * |
| **SOG [l]** | F(3,12.80) = 11.25, p = .001, pcorr = .001 | F(1,13.16) = 1.07, p = .319, pcorr = .828 | F(3,13.37) = 0.33, p = .805, pcorr = .937 | | | | | ** | * |
| **Fourth OG [l]** | F(3,15.02) = 23.00, p < .001, pcorr < .001 | F(1,14.52) = 0.61, p = .448, pcorr = .828 | F(3,13.70) = 0.25, p = .857, pcorr = .937 | *** | | | | *** | ** |
| **aIPS [l]** | F(3,15.67) = 15.13, p < .001, pcorr < .001 | F(1,16.15) = 2.50, p = .133, pcorr = .762 | F(3,15.98) = 1.54, p = .242, pcorr = .651 | *** | | | | ** | * |
| **mIPS [l]** | F(3,16.11) = 12.10, p < .001, pcorr < .001 | F(1,16.11) = 0.20, p = .659, pcorr = .828 | F(3,15.53) = 1.06, p = .395, pcorr = .747 | *** | | | | ** | * |
| **pIPS [l]** | F(3,13.14) = 5.71, p = .010, pcorr = .010 | F(1,13.19) = 0.95, p = .346, pcorr = .828 | F(3,11.85) = 1.39, p = .294, pcorr = .651 | * | | | | * | |

| | PAS effect | Emotion effect | Interaction effect | PAS pairwise comparisons | | | | | |
|---|---|---|---|---|---|---|---|---|---|
| | | | | **4>1** | **4>2** | **4>3** | **3>2** | **3>1** | **2>1** |
| **pSTS [l]** | F(3,16.00) = 7.77, p = .002, pcorr = .002 | F(1,16.40) = 0.12, p = .731, pcorr = .828 | F(3,15.17) = 1.80, p = .190, pcorr = .651 | ** | † | | | * | * |
| **LOTC [l]** | F(3,14.54) = 13.48, p < .001, pcorr < .001 | F(1,15.48) = 2.16, p = .161, pcorr = .762 | F(3,15.86) = 2.05, p = .148, pcorr = .651 | *** | * | | * | ** | |
| **IFC [l]** | F(3,16.01) = 19.50, p < .001, pcorr < .001 | F(1,9.82) = 0.42, p = .532, pcorr = .828 | F(3,13.81) = 2.11, p = .146, pcorr = .651 | *** | | | | ** | * |
| **FG [l]** | F(3,14.13) = 15.10, p < .001, pcorr < .001 | F(1,8.01) = 0.17, p = .687, pcorr = .828 | F(3,14.09) = 1.76, p = .201, pcorr = .651 | *** | | | † | *** | * |
| **ITG [l]** | F(3,13.40) = 9.65, p = .001, pcorr = .002 | F(1,10.89) = 0.26, p = .621, pcorr = .828 | F(3,12.07) = 0.01, p = .999, pcorr = .999 | ** | † | | | * | |
| **STG [l]** | F(3,7.83) = 11.03, p = .003, pcorr = .004 | F(1,14.29) = 4.11, p = .062, pcorr = .762 | F(3,9.91) = 2.88, p = .090, pcorr = .651 | * | | | | ** | * |

*Notes:* *: $p_{corrected} < .05$; **: $p_{corrected} < .01$; ***: $p_{corrected} < .001$; †: $p_{uncorrected} < .05$; †† $p_{uncorrected} < .01$; *Abbreviations*: **aIPS**: anterior intraparietal sulcus; **AMYG**: amygdala; **F**: fear; **FG**: fusiform gyrus; **IFC**: inferior frontal cortex; **ITG**: inferior temporal gyrus; **LOTC**: lateral occipito-temporal cortex; **mIPS**: medial intraparietal sulcus; **N**: neutral; **N.A.:** not applicable; **OG**: occipital gyrus; **PAS**: perceptual awareness scale; **pIPS**: posterior intraparietal sulcus; **pSTS**: posterior superior temporal sulcus; **STG**: superior temporal gyrus; **[l]**: left hemisphere; **[r]**: right hemisphere.

**Table S5. Results of comparing ROI activity against baseline for each Emotion x PAS condition.**

| | N1 | N2 | N3 | N4 | F1 | F2 | F3 | F4 |
|---|---|---|---|---|---|---|---|---|
| **ITG [r]** | t(16) = 0.89, p = .388, pcorr = .424 | t(16) = 2.57, p = .021, pcorr = .025 | t(16) = 3.35, p = .004, pcorr = .005 | t(13) = 4.87, p < .001, pcorr < .001 | t(16) = 0.88, p = .393, pcorr = .445 | t(16) = 1.69, p = .110, pcorr = .117 | t(16) = 3.37, p = .004, pcorr = .004 | t(15) = 3.45, p = .004, pcorr = .004 |
| **LOTC [r]** | t(16) = 8.25, p < .001, pcorr < .001 | t(16) = 9.62, p < .001, pcorr < .001 | t(16) = 11.66, p < .001, pcorr < .001 | t(13) = 10.09, p < .001, pcorr < .001 | t(16) = 8.29, p < .001, pcorr < .001 | t(16) = 10.07, p < .001, pcorr < .001 | t(16) = 11.18, p < .001, pcorr < .001 | t(15) = 8.55, p < .001, pcorr < .001 |
| **FG [r]** | t(16) = 8.94, p < .001, pcorr < .001 | t(16) = 9.48, p < .001, pcorr < .001 | t(16) = 11.94, p < .001, pcorr < .001 | t(13) = 9.46, p < .001, pcorr < .001 | t(16) = 9.14, p < .001, pcorr < .001 | t(16) = 9.69, p < .001, pcorr < .001 | t(16) = 10.59, p < .001, pcorr < .001 | t(15) = 9.32, p < .001, pcorr < .001 |
| **AMYG [r]** | t(16) = -0.26, p = .795, pcorr = .795 | t(16) = 2.80, p = .013, pcorr = .017 | t(16) = 2.41, p = .028, pcorr = .030 | t(13) = 3.49, p = .004, pcorr = .004 | t(16) = -0.81, p = .431, pcorr = .458 | t(16) = 1.98, p = .065, pcorr = .074 | t(16) = 3.67, p = .002, pcorr = .003 | t(15) = 3.63, p = .002, pcorr = .003 |
| **Precuneus [l]** | t(16) = 2.42, p = .028, pcorr = .036 | t(16) = 2.32, p = .034, pcorr = .036 | t(16) = 5.00, p < .001, pcorr < .001 | t(13) = 5.76, p < .001, pcorr < .001 | t(16) = 1.25, p = .229, pcorr = .278 | t(16) = 4.66, p < .001, pcorr < .001 | t(16) = 6.30, p < .001, pcorr < .001 | t(15) = 4.14, p = .001, pcorr = .001 |
| **Fourth OG [l]** | t(16) = 7.01, p < .001, pcorr < .001 | t(16) = 6.90, p < .001, pcorr < .001 | t(16) = 7.56, p < .001, pcorr < .001 | t(13) = 5.24, p < .001, pcorr < .001 | t(16) = 7.21, p < .001, pcorr < .001 | t(16) = 6.69, p < .001, pcorr < .001 | t(16) = 6.21, p < .001, pcorr < .001 | t(15) = 6.19, p < .001, pcorr < .001 |
| **SOG [l]** | t(16) = 5.47, p < .001, pcorr < .001 | t(16) = 5.19, p < .001, pcorr < .001 | t(16) = 5.31, p < .001, pcorr < .001 | t(13) = 3.84, p = .002, pcorr = .002 | t(16) = 5.92, p < .001, pcorr < .001 | t(16) = 5.41, p < .001, pcorr < .001 | t(16) = 4.87, p < .001, pcorr < .001 | t(15) = 4.66, p < .001, pcorr < .001 |
| **Fourth OG [l]** | t(16) = 8.19, p < .001, pcorr < .001 | t(16) = 7.20, p < .001, pcorr < .001 | t(16) = 7.42, p < .001, pcorr < .001 | t(13) = 4.92, p < .001, pcorr < .001 | t(16) = 8.16, p < .001, pcorr < .001 | t(16) = 6.70, p < .001, pcorr < .001 | t(16) = 6.51, p < .001, pcorr < .001 | t(15) = 6.29, p < .001, pcorr < .001 |
| **aIPS [l]** | t(16) = 4.45, p < .001, pcorr = .001 | t(16) = 5.74, p < .001, pcorr < .001 | t(16) = 5.44, p < .001, pcorr < .001 | t(13) = 5.00, p < .001, pcorr < .001 | t(16) = 4.11, p = .001, pcorr = .001 | t(16) = 5.92, p < .001, pcorr < .001 | t(16) = 6.16, p < .001, pcorr < .001 | t(15) = 6.19, p < .001, pcorr < .001 |

| | N1 | N2 | N3 | N4 | F1 | F2 | F3 | F4 |
|---|---|---|---|---|---|---|---|---|
| **mIPS [l]** | t(16) = 4.03, p = .001, pcorr = .002 | t(16) = 5.42, p < .001, pcorr < .001 | t(16) = 5.79, p < .001, pcorr < .001 | t(13) = 7.09, p < .001, pcorr < .001 | t(16) = 4.33, p = .001, pcorr = .001 | t(16) = 5.73, p < .001, pcorr < .001 | t(16) = 6.20, p < .001, pcorr < .001 | t(15) = 5.62, p < .001, pcorr < .001 |
| **pIPS [l]** | t(16) = 3.92, p = .001, pcorr = .002 | t(16) = 4.00, p = .001, pcorr = .002 | t(16) = 4.61, p < .001, pcorr < .001 | t(13) = 4.96, p < .001, pcorr < .001 | t(16) = 2.69, p = .016, pcorr = .025 | t(16) = 4.63, p < .001, pcorr < .001 | t(16) = 4.74, p < .001, pcorr < .001 | t(15) = 4.95, p < .001, pcorr < .001 |
| **pSTS [l]** | t(16) = 2.65, p = .017, pcorr = .025 | t(16) = 3.60, p = .002, pcorr = .003 | t(16) = 3.61, p = .002, pcorr = .003 | t(13) = 5.58, p < .001, pcorr < .001 | t(16) = 2.31, p = .035, pcorr = .049 | t(16) = 4.15, p = .001, pcorr = .001 | t(16) = 4.47, p < .001, pcorr = .001 | t(15) = 5.61, p < .001, pcorr < .001 |
| **LOTC [l]** | t(16) = 10.14, p < .001, pcorr < .001 | t(16) = 8.79, p < .001, pcorr < .001 | t(16) = 9.95, p < .001, pcorr < .001 | t(13) = 9.31, p < .001, pcorr < .001 | t(16) = 9.58, p < .001, pcorr < .001 | t(16) = 10.14, p < .001, pcorr < .001 | t(16) = 10.51, p < .001, pcorr < .001 | t(15) = 10.16, p < .001, pcorr < .001 |
| **IFC [l]** | t(16) = 5.59, p < .001, pcorr < .001 | t(16) = 6.51, p < .001, pcorr < .001 | t(16) = 8.34, p < .001, pcorr < .001 | t(13) = 9.46, p < .001, pcorr < .001 | t(16) = 7.39, p < .001, pcorr < .001 | t(16) = 9.63, p < .001, pcorr < .001 | t(16) = 6.80, p < .001, pcorr < .001 | t(15) = 7.90, p < .001, pcorr < .001 |
| **FG [l]** | t(16) = 7.37, p < .001, pcorr < .001 | t(16) = 8.78, p < .001, pcorr < .001 | t(16) = 10.50, p < .001, pcorr < .001 | t(13) = 9.11, p < .001, pcorr < .001 | t(16) = 6.26, p < .001, pcorr < .001 | t(16) = 8.45, p < .001, pcorr < .001 | t(16) = 9.61, p < .001, pcorr < .001 | t(15) = 8.85, p < .001, pcorr < .001 |
| **ITG [l]** | t(16) = 0.87, p = .399, pcorr = .424 | t(16) = 2.34, p = .033, pcorr = .036 | t(16) = 2.90, p = .010, pcorr = .012 | t(13) = 5.69, p < .001, pcorr < .001 | t(16) = 0.64, p = .530, pcorr = .530 | t(16) = 2.00, p = .062, pcorr = .074 | t(16) = 3.29, p = .005, pcorr = .005 | t(15) = 5.11, p < .001, pcorr < .001 |
| **STG [l]** | t(16) = 1.74, p = .102, pcorr = .123 | t(16) = -0.63, p = .539, pcorr = .539 | t(16) = -0.60, p = .557, pcorr = .557 | t(13) = 0.01, p = .989, pcorr = .989 | t(16) = 2.04, p = .058, pcorr = .075 | t(16) = -1.15, p = .268, pcorr = .268 | t(16) = -0.91, p = .379, pcorr = .379 | t(15) = -1.75, p = .101, pcorr = .101 |

***Note***: results of t-test against 0. In red, non-significant differences from baseline. *Abbreviations*: **aIPS**: anterior intraparietal sulcus; **AMYG**: amygdala; **F**: fear; **FG**: fusiform gyrus; **IFC**: inferior frontal cortex; **ITG**: inferior temporal gyrus; **LOTC**: lateral occipito-temporal cortex; **mIPS**: medial intraparietal sulcus; **N**: neutral; **N.A.:** not applicable; **OG**: occipital gyrus; **p**: p-value; **PAS**: perceptual awareness scale; **pcorr**: BHFDR corrected p-value; **pIPS**: posterior intraparietal sulcus; **pSTS**: posterior superior temporal sulcus; **SOG**: superior occipital gyrus; **STG**: superior temporal gyrus; **t**: t-value; **[l]**: left hemisphere; **[r]**: right hemisphere.

#### Table S6. Analyses on the gradual vs. dichotomous model fitting of neural data

| | Results of linear model on BIC values (Emotion x PAS) | | | Results paired t-test of gradual and dichotomous BIC model values | |
|---|---|---|---|---|---|
| | Mean ± SE | Main and interaction effects (Model, Emotion, Interaction) | Type of model | Paired-sample t-test | Type of model |
| **ITG [r]** | GN = 1.13 ± 1.14 | | | | |
| | GF = 4.01 ± 1.14 | F(1,16) = 0.20, p = .662, pcorr = .750 | | t(16) = -0.14, p = .892, | |
| | DN = 4.10 ± 1.14 | F(1,16) = 0.11, p = .745, pcorr = .907 | | pcorr > .999 | |
| | DF = 2.38 ± 1.14 | F(1,16) = 3.28, p = .089, pcorr = .504 | | | |
| **LOTC [r]** | GN = 1.04 ± 0.85 | | | | |
| | GF = 1.85 ± 0.85 | F(1,16) = 12.51, p = .003,pcorr=.047 | Gradual** | | |
| | DN = 3.80 ± 0.85 | F(1,16) = 0.63, p = .438, pcorr = .827 | | | |
| | DF = 5.67 ± 0.85 | F(1,16) = 0.27, p = .611, pcorr = .833 | | | |
| **FG [r]** | GN = 0.93 ± 0.53 | | | | |
| | GF = 2.19 ± 0.53 | F(1,16) = 3.88, p = .067, pcorr = .226 | | t(16) = -1.47, p = .161, | |
| | DN = 2.90 ± 0.53 | F(1,16) = 1.32, p = .268, pcorr = .728 | | pcorr > .999 | |
| | DF = 4.81 ± 0.53 | F(1,16) = 0.08, p = .784, pcorr = .833 | | | |
| **AMYG** | GN = 0.91 ± 0.63 | | | | |
| **[r]** | GF = 3.84 ± 0.63 | F(1,16) = 0.10, p = .760, pcorr = .808 | | t(16) = -0.19, p = .853, | |
| | DN = 1.59 ± 0.63 | F(1,16) = 1.77, p = .202, pcorr = .728 | | pcorr > .999 | |
| | DF = 3.60 ± 0.63 | F(1,16) = 0.24, p = .634, pcorr = .833 | | | |
| **Precuneus** | GN = 4.00 ± 1.57 | | | | |
| **[l]** | GF = 4.82 ± 1.57 | F(1,16) = 0.28, p = .606, pcorr = .750 | | t(16) = -0.35, p = .730, | |
| | DN = 4.89 ± 1.57 | F(1,16) = 0.07, p = .790, pcorr = .907 | | pcorr > .999 | |
| | DF = 5.08 ± 1.57 | F(1,16) = 0.36, p = .557, pcorr = .833 | | | |
| **Fourth** | GN = 2.71 ± 1.02 | | | | |
| **OG [l]** | GF = 5.99 ± 1.02 | F(1,16) = 0.56, p = .467, pcorr = .750 | | t(16) = -1.37, p = .189, | |
| | DN = 4.74 ± 1.02 | F(1,16) = 0.97, p = .340, pcorr = .728 | | pcorr > .999 | |
| | DF = 5.50 ± 1.02 | F(1,16) = 0.89, p = .359, pcorr = .833 | | | |
| **SOG [l]** | GN = 0.18 ± 1.03 | | | | |
| | GF = 3.45 ± 1.03 | F(1,16) = 5.62, p = .031, pcorr = .211 | N-Gradual** | | |
| | DN = 4.78 ± 1.03 | F(1,16) = 0.10, p = .754, pcorr = .907 | F-Dichotomous | | |
| | DF = 3.21 ± 1.03 | F(1,16) = 6.73, p = .020, pcorr = .332 | | | |
| **Fourth** | GN = 2.69 ± 0.75 | | | | |
| **OG [l]** | GF = 0.40 ± 0.75 | F(1,16) = 0.21, p = .655, pcorr = .750 | | t(16) = 0.86, p = .404, | |
| | DN = 3.87 ± 0.75 | F(1,16) = 2.73, p = .118, pcorr = .728 | | pcorr > .999 | |
| | DF = 0.14 ± 0.75 | F(1,16) = 0.93, p = .348, pcorr = .833 | | | |
| **aIPS [l]** | GN = 1.95 ± 1.57 | | | | |
| | GF = 3.17 ± 1.57 | F(1,16) = 0.33, p = .573, pcorr = .750 | N-Dichotomous | | |
| | DN = -0.04 ± 1.57 | F(1,16) = 4.31, p = .054, pcorr = .728 | F-Gradual | | |
| | DF = 3.81 ± 1.57 | F(1,16) = 4.50, p = .050, pcorr = .424 | | | |
| **mIPS [l]** | GN = 2.99 ± 1.66 | | | | |
| | GF = 2.48 ± 1.66 | F(1,16) = 5.16, p = .037, pcorr = .211 | Dichotomous* | | |
| | DN = 0.58 ± 1.66 | F(1,16) = 0.01, p = .943, pcorr = .943 | | | |
| | DF = 0.79 ± 1.66 | F(1,16) = 0.10, p = .751, pcorr = .833 | | | |
| **pIPS [l]** | GN = 2.72 ± 0.65 | | | | |
| | GF = 4.36 ± 0.65 | F(1,16) = 3.89, p = .066, pcorr = .226 | | t(16) = 1.33, p = .201, | |
| | DN = -0.10 ± 0.65 | F(1,16) = 1.79, p = .199, pcorr = .728 | | pcorr > .999 | |
| | DF = 2.14 ± 0.65 | F(1,16) = 0.11, p = .747, pcorr = .833 | | | |
| **pSTS [l]** | GN = -0.20 ± 0.75 | | | | |
| | GF = 1.58 ± 0.75 | F(1,16) = 1.81, p = .197, pcorr = .420 | | t(16) = -0.07, p = .947, | |
| | DN = 1.94 ± 0.75 | F(1,16) = 0.96, p = .342, pcorr = .728 | | pcorr > .999 | |
| | DF = 3.06 ± 0.75 | F(1,16) = 0.10, p = .753, pcorr = .833 | | | |
| **LOTC [l]** | GN = 2.31 ± 0.49 | F(1,16) = 2.33, p = .146, pcorr = .356 | | t(16) = -1.00, p = .334, | |
| | GF = 3.23 ± 0.49 | F(1,16) = 0.02, p = .896, pcorr = .943 | | pcorr > .999 | |
| | DN = 4.55 ± 0.49 | F(1,16) = 0.28, p = .605, pcorr = .833 | | | |
| | DF = 4.06 ± 0.49 | | | | |
| **IFC [l]** | GN = 3.48 ± 1.57 | | | | |
| | GF = 5.27 ± 1.57 | F(1,16) = 0.96, p = .341, pcorr = .644 | | t(16) = 2.14, p = .048, | |
| | DN = 2.74 ± 1.57 | F(1,16) = 0.46, p = .506, pcorr = .860 | | pcorr > .999 | Dichotomous* |
| | DF = 2.79 ± 1.57 | F(1,16) = 0.42, p = .528, pcorr = .833 | | | |

| | Mean ± SE | Main and interaction effects (Model, Emotion, Interaction) | Type of model | Paired-sample t-test | Type of model |
|---|---|---|---|---|---|
| **FG [l]** | GN = 0.95 ± 0.69 | F(1,16) = 0.01, p = .942, pcorr = .942 | | t(16) = 1.02, p = .324, | |
| | GF = 2.90 ± 0.69 | F(1,16) = 0.11, p = .742, pcorr = .907 | | pcorr > .999 | |
| | DN = 2.15 ± 0.69 | F(1,16) = 2.34, p = .145, pcorr = .618 | | | |
| | DF = 1.49 ± 0.69 | | | | |
| **ITG [l]** | GN = 3.49 ± 1.26 | | | | |
| | GF = 4.42 ± 1.26 | F(1,16) = 0.39, p = .540, pcorr = .750 | | t(16) = 0.39, p = .698, | |
| | DN = 3.41 ± 1.26 | F(1,16) = 0.07, p = .800, pcorr = .907 | | pcorr > .999 | |
| | DF = 3.34 ± 1.26 | F(1,16) = 0.86, p = .368, pcorr = .833 | | | |
| **STG [l]** | GN = 1.74 ± 0.36 | | | | |
| | GF = 3.41 ± 0.36 | F(1,16) = 2.83, p = .112, pcorr = .318 | | t(16) = -1.32, p = .204, | |
| | DN = 3.08 ± 0.36 | F(1,16) = 1.12, p = .305, pcorr = .728 | | pcorr > .999 | |
| | DF = 4.84 ± 0.36 | F(1,16) = 0.00, p = .953, pcorr = .953 | | | |

*Abbreviations*: *: **p** < .05; **aIPS**: anterior intraparietal sulcus; **AMYG**: amygdala; **BIC:** Bayesian Information Criterion; **DN:** dichotomous model for neutral bodies; **DF:** dichotomous model for fearful bodies; **F**: fear; **FG**: fusiform gyrus; **GN:** gradual model for neutral bodies; **GF:** gradual model for fearful bodies; **IFC**: inferior frontal cortex; **ITG**: inferior temporal gyrus; **LOTC**: lateral occipito-temporal cortex; **mIPS**: medial intraparietal sulcus; **N**: neutral; **N.A.:** not applicable; **OG**: occipital gyrus; **p**: p-value; **PAS**: perceptual awareness scale; **pcorr**: BHFDR corrected p-value; **pIPS**: posterior intraparietal sulcus; **pSTS**: posterior superior temporal sulcus; **SE:** standard error; **SOG**: superior occipital gyrus; **STG**: superior temporal gyrus; **t**: t-value; **[l]**: left hemisphere; **[r]**: right hemisphere.

**Table S7. Analysis of model slopes and intercepts of neural data**

| | Results paired t-test of Neutral vs. Fear model slopes comparison | | Results paired t-test of Neutral vs. Fear model intercept comparison | |
|---|---|---|---|---|
| | **Mean ± SE** | **Paired-sample t-test** | **Mean ± SE** | **Paired-sample t-test** |
| **LOTC [r]** | N = 1.38 ± 0.24<br>F = 0.89 ± 0.20 | t(16) = 1.51, p = .150,<br>pcorr > .999 | N = 1.44 ± 0.18<br>F = 1.54 ± 0.17 | t(16) = -0.67, p = .514,<br>pcorr > .999 |
| **mIPS [l]** | N = 0.72 ± 0.15<br>F = 0.71 ± 0.17 | t(16) = 0.05, p = .959,<br>pcorr > .999 | N = 0.79 ± 0.19<br>F = 0.79 ± 0.21 | t(16) = 0.00, p = .999,<br>pcorr > .999 |
| **IFC [l]** | N = 1.06 ± 0.17<br>F = 0.78 ± 0.21 | t(16) = 1.40, p = .179,<br>pcorr > .999 | N = 0.75 ± 0.13<br>F = 0.93 ± 0.13 | t(16) = -1.31, p = .207,<br>pcorr > .999 |

*Abbreviations*: **F**: fear; **IFC**: inferior frontal cortex; **LOTC**: lateral occipito-temporal cortex; **mIPS**: medial intraparietal sulcus; **N**: neutral; **p**: p-value; **PAS**: perceptual awareness scale; **pcorr**: BHFDR corrected p-value; **SE:** standard error; **t**: t-value; **[l]**: left hemisphere; **[r]**: right hemisphere.

**Supplementary information references**

Alvarez, I., de Haas, B., Clark, C. A., Rees, G., & Schwarzkopf, D. S. (2015). Comparing different stimulus configurations for population receptive field mapping in human fMRI. Frontiers in human neuroscience, 9, 96. doi:10.3389/fnhum.2015.00096

Brainard, D. H., & Vision, S. (1997). The psychophysics toolbox. Spatial vision, 10(4), 433-436. doi:10.1163/156856897X00357

de Gelder, B., & Van den Stock, J. (2011). The bodily expressive action stimulus test (BEAST). Construction and validation of a stimulus basis for measuring perception of whole body expression of emotions. Frontiers in psychology, 2, 181. doi:10.3389/fpsyg.2011.00181

Langner, O., Dotsch, R., Bijlstra, G., Wigboldus, D. H., Hawk, S. T., & Van Knippenberg, A. (2010). Presentation and validation of the Radboud Faces Database. Cognition and emotion, 24(8), 1377-1388. doi:10.1080/02699930903485076

Lundqvist, D., Flykt, A., & Öhman, A. (1998). Karolinska directed emotional faces. Cognition and emotion. doi:10.1037/t27732-000

Peirce, J. W. (2007). PsychoPy—psychophysics software in Python. Journal of neuroscience methods, 162(1-2), 8-13. doi:10.1016/j.jneumeth.2006.11.017

Peirce, J. W. (2009). Generating stimuli for neuroscience using PsychoPy. Frontiers in neuroinformatics, 2, 10. doi:10.3389/neuro.11.010.2008

Pelli, D. G., & Vision, S. (1997). The VideoToolbox software for visual psychophysics: Transforming numbers into movies. Spatial vision, 10, 437-442. doi:10.1163/156856897X00366

Schwarzkopf, D. S., Anderson, E. J., de Haas, B., White, S. J., & Rees, G. (2014). Larger extrastriate population receptive fields in autism spectrum disorders. Journal of neuroscience, 34(7), 2713-2724. doi:10.1523/JNEUROSCI.4416-13.2014

van Dijk, J. A., de Haas, B., Moutsiana, C., & Schwarzkopf, D. S. (2016). Intersession reliability of population receptive field estimates. Neuroimage, 143, 293-303.
